# Molecular Noise-Filtering in the *β*-adrenergic Signaling Network by Phospholamban Pentamers

**DOI:** 10.1101/2020.12.30.424828

**Authors:** Daniel Koch, Alexander Alexandrovich, Florian Funk, Joachim P. Schmitt, Mathias Gautel

## Abstract

Phospholamban (PLN) is an important regulator of calcium handling in cardiomyocytes due to its ability to inhibit the sarco(endo)plasmic reticulum calcium-ATPase (SERCA) *β*-adrenergic stimulation reverses SERCA inhibition via PLN phosphorylation and facilitates fast calcium reuptake PLN also forms pentamers whose physiological significance has remained elusive Using biochemical experiments and mathematical modeling, we show that pentamers regulate both the dynamics and steady-state levels of monomer phosphorylation Substrate competition by pentamers and a feed-forward loop involving inhibitor-can delay monomer phosphorylation by protein kinase A (PKA) Steady-state phosphorylation of PLN is predicted to be bistable due to cooperative dephosphorylation of pentamers Both effects act as complementary noise-filters which can reduce the effect of random fluctuations in PKA activity Pentamers thereby ensure consistent monomer phosphorylation and SERCA activity in spite of noisy upstream signals Preliminary analyses suggest that the PLN mutation R del could impair noise-filtering, offering a new perspective on how this mutation causes cardiac arrhythmias.

## 1 Introduction

Calcium (Ca^2+^) currents determine contraction and relaxation of the heart at the cellular level: high C^2+^ concentrations enable sarcomeric cross-bridge cycling leading to contraction, low Ca ^+^ concentrations lead to relaxation[1]These currents are controlled by the release and reuptake of calcium from and into the sarcoplasmic reticulum (SR), the major storage compartment for intracellular Ca^2+^ At the molecular level, dozens of proteins regulate Ca^2+^ -handling and excitation-contraction coupling [3] The Ca^2+^ pump SERCA mediates approximately 0-0% of the Ca^2+^ reuptake into the SR and therefore induces relaxation of the cardiomyocyte [1, 4] SERCA function is inhibited by phospholamban (PLN), a 52 amino-acid protein resident in the SR membrane Phosphorylation of PLN at Ser by protein kinase A (PKA) reverses SERCA inhibition in response to *β*-adrenergic stimulation and thereby accelerates Ca^2+^ removal and cardiomyocyte relaxation [4-8] This constitutes an important mechanism to adapt cardiac output to increasing demand and is an integral part of the *β*-adrenergic “fight-or-flight” response [4, 8,9] Disruptions in this part of the *β*-adrenergic signaling network can have drastic consequences multiple mutations in the PLN gene have been discovered in the past two decades, most of which cause severe forms of cardiomyopathy leading to cardiac arrhythmias and heart failure[1 0-14].

In spite of the progress in understanding the structure and function of PLN, many aspects of this protein are still poorly understood and specific therapeutic approaches to manipulate the PLN signaling network are lacking One of the less well understood aspects is the assembly of PLN into homo-pentamers[15] Although their pinwheel-like structure in lipid environments [16] yields intuitive plausibility to early conjectures and data suggesting that pentameric PLN acts as an ion channel [17, 8], this hypothesis has been contested by multiple experimental, structural and theoretical studies [19 - 21] Since an artificial monomeric PLN mutant was found to be a similarly potent SERCA inhibitor as wild-type PLN [22], the prevailing paradigm considers pentamers to be a biologically inactive storage form[4,8,20] owever, increasing evidence suggests that PLN pentamers are not entirely passive and influence cardiomyocyte contractility and PLN phosphorylation dynamics [23-26] ostrikov proposed that pentamers could act as buffers which fine-tune SERCA regulation via monomeric PLN by keeping it within a physiological window [20] Yet, it is not obvious what benefit exactly pentamerization contributes given that SERCA activity can already be controlled by regulating expression levels and by multiple post-translational modifications of both PLN and SERCA [4,27,28] The specific physiological advantage of pentamerization and its role in the pathophysiology of PLN mutations remains thus elusive.

In the present study, we investigated the role of pentamers in the PLN regulatory network We found that pentamers have only a limited capacity to buffer the concentration of monomeric phospholamban since the effect is slow and moderate Based on the hypothesis that the function of pentameric PLN exceeds monomer buffering, we developed a mathematical model of the PLN regulatory network to study the role of pentamers in the context of *β*-adrenergic stimulation from a dynamical systems perspective Our results indicate that pentamers are molecular noise-filters to ensure consistent PLN phosphorylation in response to noisy *β*-adrenergic stimulation A preliminary analysis of the arrhythmogenic PLN mutation R del suggests that this mutation could impair noise-filtering, indicating that molecular noise-filtering in the *β*-adrenergic signaling network could be important to prevent cardiac arrhythmias.

## 2 Results

### 2.1 Pentamers are moderate and slow monomer buffers *in vitro*

The predominant paradigm is that PLN pentamers are a storage or buffering reservoir for monomers (Figure 1A), [4,8, 20, 21] The oligomeric state of PLN in tissue or cell homogenates is typically assessed from samples in SDS sample buffer which does not interfere with oligomerization owever, SDS is a harsh anionic detergent which interferes with the function of many other proteins We therefore studied PLN oligomerization in both SDS sample buffer and a Triton™ X-100 based buffer (TBB) at physiological p and ionic strength which effectively solubilises PLN and allows for rapid phosphorylation of PLN at S by PKA.

To test the hypothesis that PLN pentamers buffer monomer concentration, we analysed the oligomeric state of PLN (unphosphorylated and phosphorylated) by semi-native SDS-PAGE at various total PLN concentrations after dilution and two hours equilibration (Figure 1B) As shown in Figure 1C, the slope of pentameric PLN in TBB is steeper than the slope of monomeric PLN (particularly at low total concentrations), suggesting that changes in total PLN concentration have a larger effect on pentameric than on monomeric PLN In contrast to the TBB samples, the slope of both pentameric and monomeric PLN in SDS sample buffer does not change much upon dilution, possibly indicating incomplete dissociation of pentamers in SDS In the likely region of physiological PLN concentrations at the SR (>5 0 µ ; Table S1), the change in monomers relative to the change in total PLN is slightly lower than for pentamers, indicating that pentamers can indeed buffer the concentration of monomers (Figure 1C, inset) However, we found that pentamers dissociate only slowly with an apparent mean lifetime (*k*_*obs*_)^−1^ of 11.4 of minutes for pentamers (Figure S1), in good agreement with previous live-cell measurements[29] We concluded that under the investigated *in vitro* conditions, PLN pentamers buffer the concentration of PLN monomers only moderately and slowly We thus hypothesized that PLN pentamers might play further roles.

**Figure 1.**
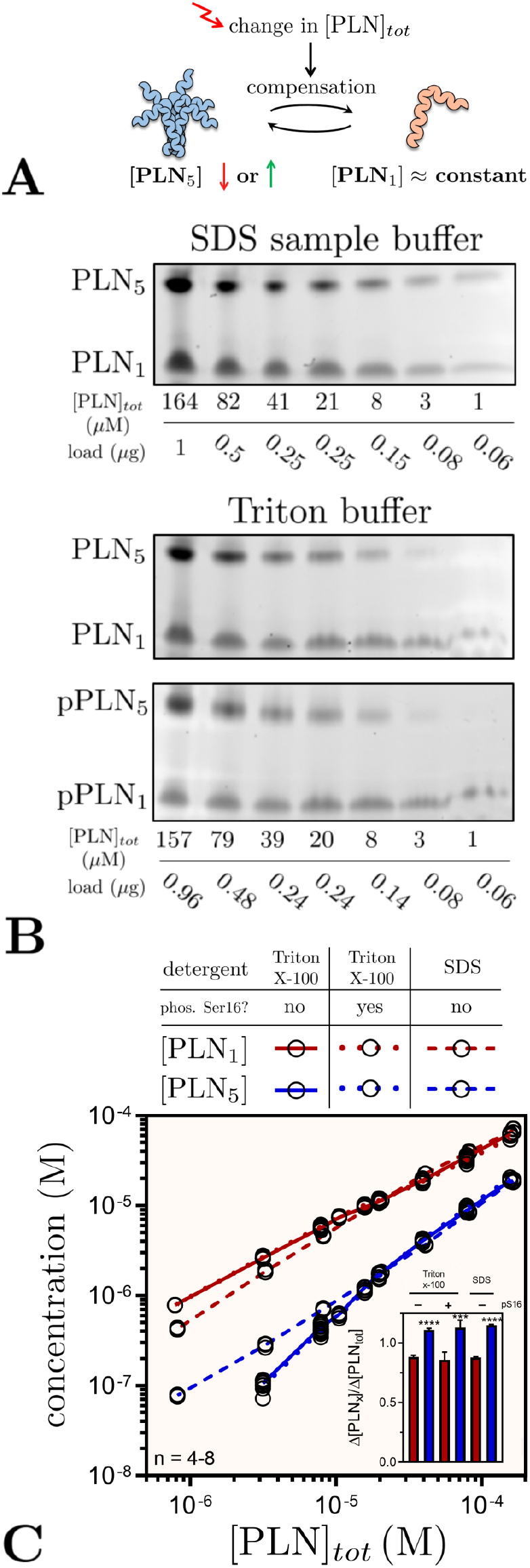
(A) In the prevailing paradigm pentamers buffer the PLN monomer concentration by compensating changes via association or dissociation (B,C) Oligomeric state of PLN in sample or Triton™ X-based buffer (TBB) at different total protein concentrations after dilution. (B)Oriole™ stained gels after semi-native SDS-PAGE (C) Quantified monomers and pentamers at different total concentrations Inset: relative change in monomers and pentamers when comparing highest two total concentrations ***p<0.001,****p<0.0001 vs monomers.

On a site note, we observed no increased pentamerization upon phosphorylation of PLN in TBB (Figure 1B,C and S2). The increase in pentamerization upon phosphorylation is sometimes called the *dynamic equilibrium* of PLN and while its biological significance is unclear, it has been speculated that it might contribute to SERCA regulation [8, 30, 31] Interestingly, we observed a significant increase in pentamerization after diluting PLN phosphorylated in TBB with SDS-sample buffer (Figure S19C), indicating that the effect relies on anionic environments.

#### 2.2 A mathematical model of the PLN regulatory network

Mathematical modeling has been paramount to understand the non-linear behavior of signaling networks and how they regulate cellular activities including growth, differentiation, apoptosis and motility PLN, too, is part of a complex signaling network involving multple kinases, phosphatases and regulatory complexes; a network which so far remained largely unexplored by mathematical approaches Although a PLN submodule is part of several models of cardiac Ca^2 +^–cycling[32] or *β*-adrenergic signal transduction [33], no mathematical model has, to our knowledge, considered PLN pentamers or provided a detailed analysis of the network immediately implicated in regulating PLN Aiming to fill this gap we set out to develop a mathematical model of the PLN network to study its functionality and the role of pentamers in the context of *β*-adrenergic stimulation from a dynamical systems perspective.

We began model development by considering several possible models of how PLN forms pentamers and calibrated them using our dilution and dissociation time-course data We found that a model following a monomer → dimer→ tetramer→ pentamer pathway shows good agreement with our experimental data and outperforms other model variants (supplementary section 2 and Figure S12).

We extended the PLN oligomerization model by including key proteins and reactions of the *β*-adrenergic signal transduction network involved in regulating Ser phosphorylation of PLN We accounted for reactions and enzymes responsible for addition (PKA) and removal of the Ser16 phosphate group (phosphatases PP1 and PP2A), [4,34] Dephosphorylation of PLN pentamers has been shown to exhibit strong positive cooperativity [35] Since PP is the main phosphatase for reversing S phosphorylation of PLN [34,36], we assumed that the catalytic turnover for dephosphorylation of pentameric PLN by PP1 increases with fewer phosphate groups left on a pentamer We implemented this assumption by introducing dimensionless parameters *φ* and *χ* for tuning individual steps of pentamer dephosphorylation by PP (Figure S16) We also included regulation of PP by inhibitor-as described in[33] Inhibitor-1 can bind and inhibit PP1 when phosphorylated by PKA at Thr35, whereas phosphorylation at this site is reversed by PP A [8,33] To keep our analysis focussed on the regulation of PLN phosphorylation in the context of *β*-adrenergic stimulation, we treated the concentration of active PKA at the SR as a model input parameter and omitted processes upstream of PKA (such as cAMP production and degradation) and downstream of PLN (such as SERCA activity and Ca^2+^-handling). Due to lack of mechanistic and kinetic data, we did not include (de-)phosphorylation of PLN at Ser10 or Thr17.

Figure 2 shows a simplified scheme of the biochemical reactions included in our model. The model comprises 60 biochemical reactions between 20 molecular species which are described by a set of 17 ordinary differential and 3 algebraic equations. The additional protein concentrations and model parameters not determined by our own data are based on experimental measurements from the literature A more detailed description of how the model was formulated can be found in the supplementary material along with the model equations and parameter values Having a mathematical description of the processes which regulate PLN phosphorylation at our disposal, we set out to explore the behavior of our model.

**Figure 2.**
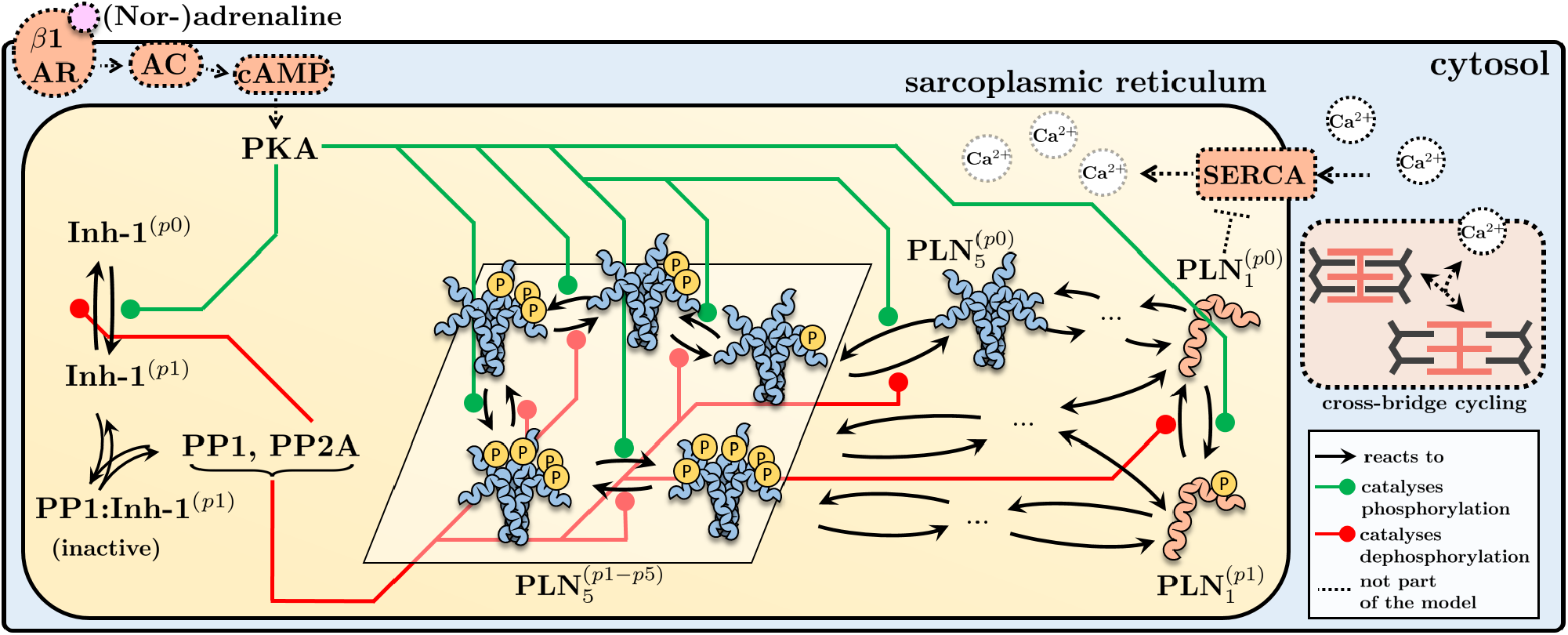
Simplified model scheme of the PLN signaling network involved in the control of SERACA activity in response to *β*-adrenergic stimulation The model captures all processes immediately involved in regulating PLN phosphorylation at Ser16 To simplify the scheme the different oligomerization routes of phosphorylated and unphosphorylated PLN are not shown See supplementary section 2 for the complete model scheme parameter values and model equations.

### 2.3 Pentamers and the inhibitor-1 feed-forward loop delay monomer phosphorylation

In a first simulation, we studied the dynamics of PLN monomer and pentamer phosphorylation by PKA in the absence of phosphatases (Figure 3A, left) The phosphorylation of monomers resembles a hyperbola but features a kink in the middle Pentamer phosphorylation, on the other hand, exhibits dynamics typical for multisite phosphorylation systems with transient waves of incompletely phosphorylated intermediate forms Next, we simulated dephosphorylation of completely phosphorylated PLN in the absence of PKA (Figure 3A, right) Expectedly, dephosphorylation resembles the phosphorylation dynamics but in reverse order As expected from the implemented cooperativity of PP1, the accumulation of unphosphorylated pentamers is more abrupt.

**Figure 3.**
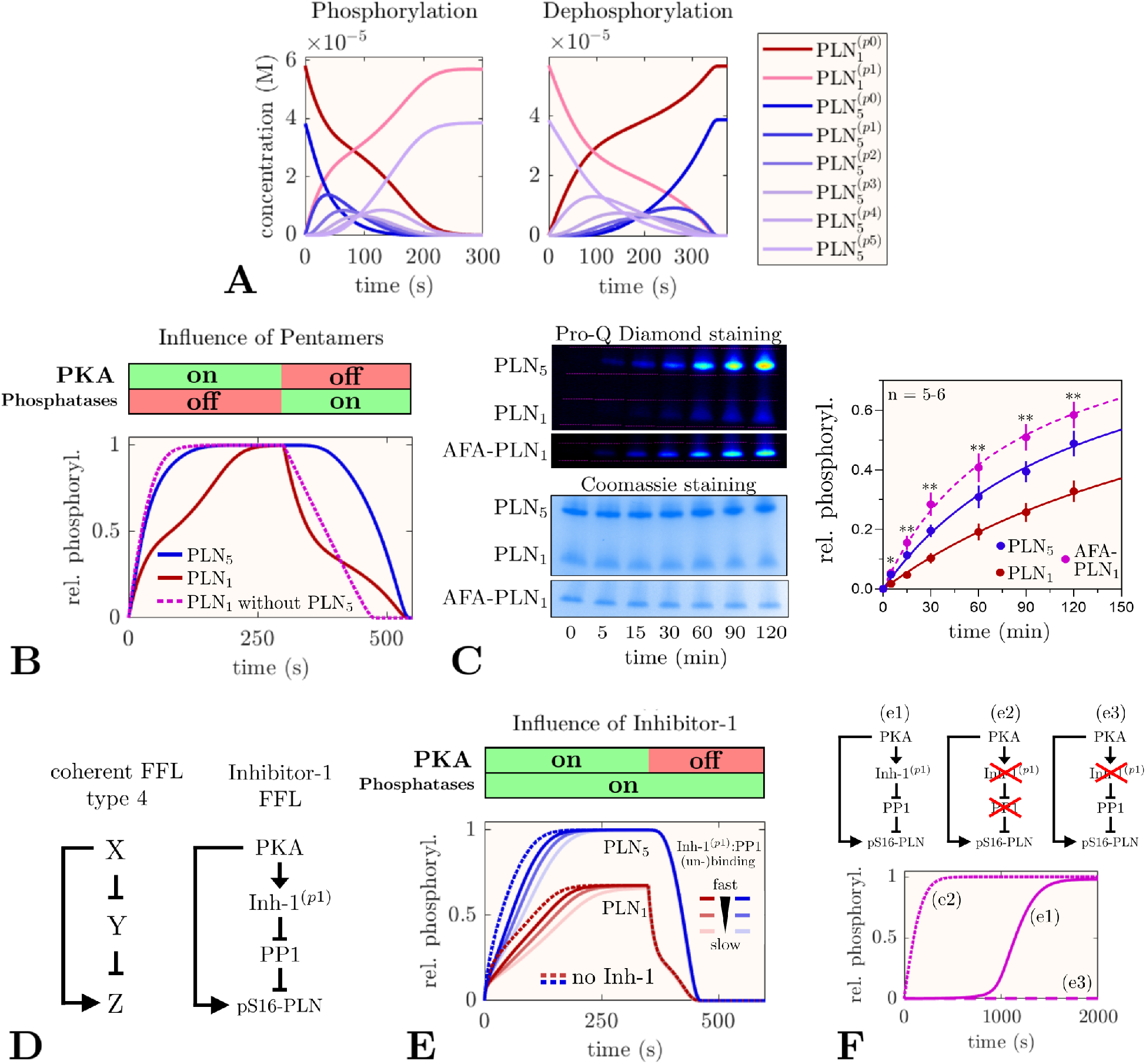
Dynamic regulation of PLN phosphorylation. (A) Time-course simulations of PLN phosphorylation by 0.1 *µM* PKA in the absence of phosphatases (left) and dephosphorylation of completely phosphorylated PLN by PP1 and PP2A in the absence of PKA (right). (B) Dynamics of relative phosphorylation levels in a sequential phosphorylation/dephosphorylation simulation in the presence or absence of pentameric PLN. (C) Experimental phosphorylation time-course of wild-type PLN ([PLN_*tot*_] ≈57 *µ*M at which [PLN_1_] ≈52 *µ*M) and monomeric AFA-PLN (≈52 *µ*M) A low PKA concentration of 25 U/*µ*L (≈7.7 nM) was chosen to slow down the reaction for easier sampling .Data represent mean ± SEM *p<0.05,**p<0.01, AFA-PLN_1_ vs PLN_1_. (D) Left: coherent FFL type 4 in which the full response of Z is delayed until the inhibitory effect of Y is revoked by X. Right: structure of the inhibitor-1 FFL.(E) Influence of the inhibitor-1 FFL in simulations of PLN phosphorylation by PKA in the presence of PP1. (F) Optimal experimental design and controls for detecting PLN phosphorylation delay by the inhibitor-1 FFL is given by [PKA] < [PP1] < [Inh-1] and indicated ‘ knock -out’ versions of the FFL.

To simplify the plots we decided to focus on relative PLN phosphorylation for the remainder of this study Interestingly, replotting the data from the phosphorylation time-course simulations reveals that relative phosphorylation of monomers significantly lags behind relative phosphorylation of pentamers (Figure 3B) A likely explanation for this delay could be that monomers and pentamers compete against each other as PKA-substrates Performing the same simulation without pentamers but at equimolar monomer concentration abolishes delayed phosphorylation, confirming that the lag is indeed caused by competing PLN pentamers (Figure 3B, dotted line) Parameters which increase substrate competition or, surprisingly, slow down pentamer phosphorylation, can increase this delay (Figure S3).

To test the predicted delay experimentally, we carried out PKA-phosphorylation time-course experiments using wild-type PLN and AFA-PLN (an artificial monomeric mutant) at equimolar monomer concentrations In agreement with the simulations, we found monomer phosphorylation to be significantly delayed in the presence of pentamers (Figure 3C).

Substrate competition is not the only network motif able to delay the response to a stimulus Interestingly, the PLN network contains a second motif with such ability: the inhibition of PP by PKA via phosphorylation of inhibitor-constitutes a subgraph which can be described as an elongated version of a coherent type feed-forward loop (FEL) able to cause delays (Figure 3D) [37] Simulations show that inhibitor-1 can indeed delay phosphorylation of PLN monomers and pentamers if binding of phosphorylated inhibitor-1 to PP1 is not too fast (Figure 3E) Reducing the PP1 concentration by the fraction which is inhibited by inhibitor-1 at steady state and repeating the simulation in the absence of inhibitor-shows that PLN phosphorylation approaches the same steady-state levels but much faster (Figure 3E, dotted lines) or slower inhibitor-phosphorylation, the delay becomes more pronounced (Figure S4) The delay can be uncoupled from pentamer competition by using monomeric AFA-PLN and maximized when [PKA] < [PP1] < [Inh-1] When contrasted to ‘knock-out’ variants of the FFL, this yields an optimal design for future experimental testing of the predicted delay (Figure 3F).

In summary, our simulations predict the existence of two independent response delay elements in the PLN network: pentamers delaying the phosphorylation of monomers and an inhibitor-L delaying the phosphorylation of both monomers and pentamers.

### 2.4 Bistability in the steady-state phosphorylation of PLN

Mangan and Alon proposed that response delay elements may act as persistence sensors which reject short input stimuli [37] Before exploring what the physiological advantage of such persistence sensing in the context of *β*-adrenergic stimulation might be, we shall first consider how PLN phosphorylation is controlled at steady state.

Multisite phosphorylation systems can exhibit ultrasensitivity and bistability if there is sufficient kinetic asymmetry in the subsequent cycles of phosphorylation and dephosphorylation e g due to cooperativity or multi-enzyme regulation [38,39] Since cooperativity is present in the dephosphorylation of pentameric PLN[35], we wondered whether PLN phosphorylation might be bistable at some level of PKA activity A hallmark of bistability is that the approached steady state depends on the system’s history (hysteresis) We therefore performed several simulations with identical conditions except for different initial levels of relative phosphorylation and found that PLN phosphorylation is indeed bistable at some PKA concentrations (Figure 4A) To better understand how steady-state PLN phosphorylation depends on PKA concentration, we performed a bifurcation analysis and found that PLN phosphorylation increases in a switch-like, ultrasensitive fashion as it passes a threshold at about 1/3of maximum PKA concentration at the SR (≈0.6 *µ*M [33]) (Figure 4B).

**Figure 4.**
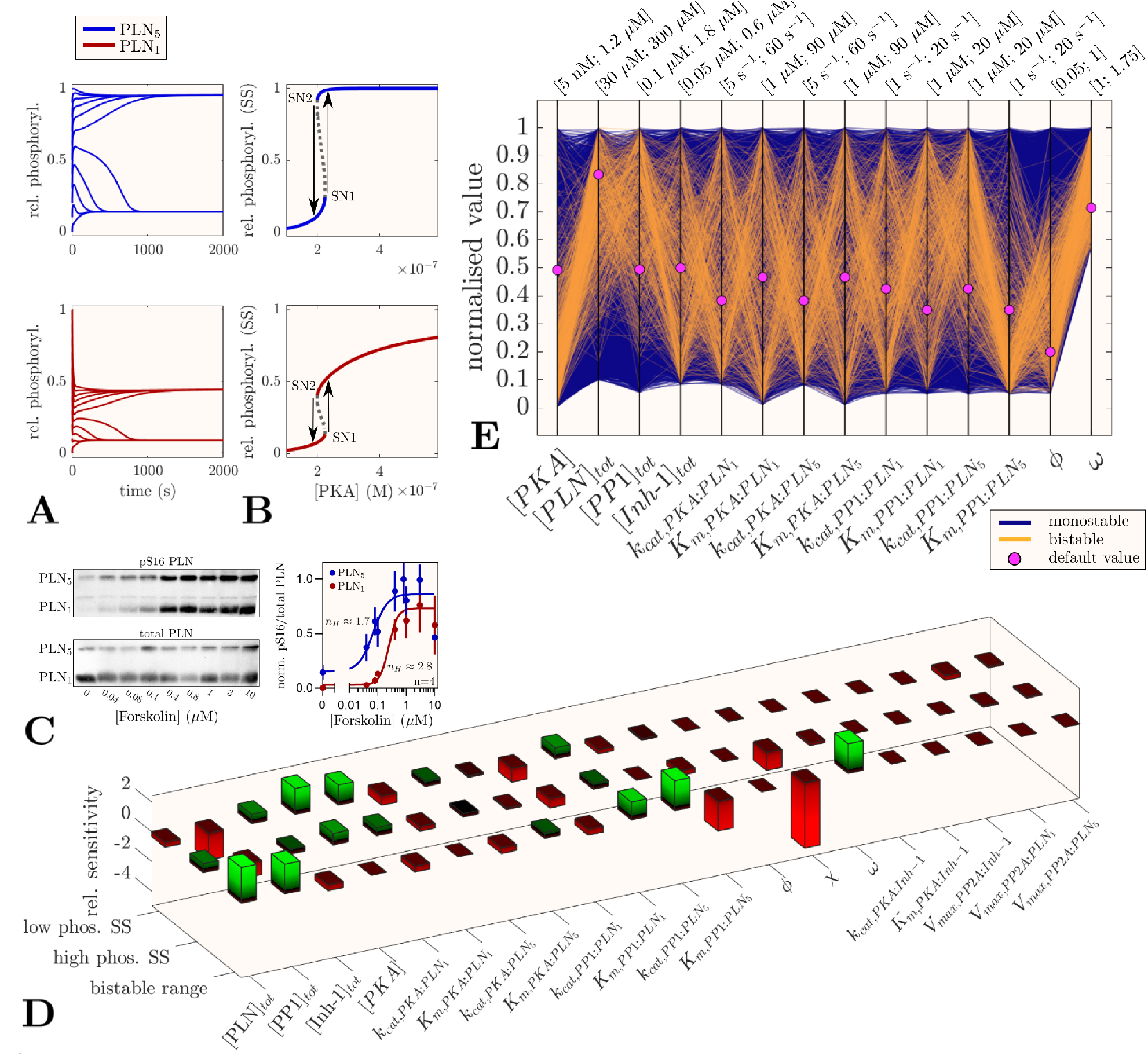
PLN phosphorylation at steady state (A) Time-course simulations with different initial levels of PLN phosphorylation show hysteresis ([PKA] 2 *µ*M) (B) Bifurcation diagrams for relative phosphorylation of PLN monomers and pentamers show a bistable region of ≈ 25 *µ*M range (C) PKA-dependent PLN phosphorylation in HEK293 cells ata represent mean ± SEM (D) Local sensitivity analysis of low phosphorylation steady state ([PKA]=0.13 *µ*M) high phosphorylation steady state ([PKA] = 0. 25 *µ*M) and of the range of the bistable region elative sensitivities determined at Δ*p=+*1*%* for high/low phosphorylation steady states and Δ;*p= +*10% for bistable range (E) Parallel coordinate plot of the model stability behavior for random parameter sets.

Since ultrasensitivity is considered a prerequisite and indication for bistability, we experimentally determined the dose-response of PLN S16 -phosphorylation in transfected HEK cells after PKA activation by forskolin and obtained fitted Hill-exponents of 1.7 for pentamers and 2.8 for monomers (Figure 4C) This confirms that the network is capable of ultrasensitive PLN phosphorylation.

To find out which model parameters exert most control over PLN phosphorylation at steady state, we performed a local sensitivity analysis of relative monomer phosphorylation (Figure 4D).We found that the low phosphorylation state is generally more sensitive to parameter perturbations than the high phosphorylation state, but both are primarily controlled by concentrations and catalytic constants of PKA and PP1. Parameters associated with inhibitor-1 or PP2A show only minor influence The bistable range depends primarily on parameters which influence PP1 cooperativity or substrate competition between PLN monomers and pentamers, e.g. higher concentrations of PLN and PP1 or changes to PP1 dependent dephosphorylation Interestingly, cooperative increase of substrate affinity (*χ*) and the dynamic equilibrium of PLN (*ω*) show a strong influence on the bistable range By default, we assumed PLN turnover (*k*_*cat*_) rather than substrate affinity (*K*_*m*_) to be regulated cooperatively (i.e. *χ =* and *φ <* 1), but the exact nature of cooperative PLN pentamer dephosphorylation is currently unknown While cooperative increase of *k*_*cat*_ is essential for the emergence of bistability, increasing substrate affinity appears to reduce the bistable range To better understand how the control parameters *φ, χ* and *ω* shape the PLN phosphorylation response curve, we performed multiple bifurcation analyses Although low *φ* and *χ* or high *ω* values can all increase the bistable range, the parameters differ markedly in how they shape other characteristics of the dose-response curve, possibly due to distinct effects on dephosphorylation rates (Figure S5).

Local sensitivity analysis allows to study the influence of parameters only around a nominal steady state, limiting the generality of its conclusions, whereas bifurcation analysis can be challenging and is limited to varying only few parameters simultaneously We therefore implemented a recently developed method which allows exploring models in a fashion unbiased by a particular parameter set by simultaneously probing an arbitrary subset of the multi-dimensional parameter space and visualizing the resulting stability behavior on parallel coordinate plots[40]. For each analysis we probed 10000 randomly sampled parameter sets focussing on the concentrations of PKA, PP1, PLN and inhibitor-1, enzymatic constants, cooperativity parameters *φ, χ* and the dynamic equilibrium of PLN (*ω*) In the absence of cooperative substrate affinity of PLN dephosphorylation (*χ*), % of the sampled parameter sets led to bistable phosphorylation responses The emergence of bistability is favored by high *k*_*cat*_ and low *K*_*m*_ values for pentamer dephosphorylation by PP1 (Figure 4E) In contrast, other PKA and PP1 constants show relatively little influence The emergence of bistability is further associated with low [PKA], high [PLN]_*tot*_ and [PP1]_*tot*_ as well as strong cooperative increase of *k*_*cat*_ of PP1 (low *φ* values) and strong dynamic equilibrium (high *ω* values) To further study the role of pentameric PLN and the nature of PP1 cooperativity in pentamer dephosphorylation, we repeated the analysis without pentameric PLN and with cooperative substrate affinity of PLN dephosphorylation (*χ* > 1), respectively We found no bistability in the absence of pentameric PLN and markedly fewer (1.1 %) parameter sets leading to bistability when *χ* > 1 (Figure S6).

In summary, these analyses show that pentamers, their cooperative dephosphorylation and the dynamic equilibrium of PLN are important factors in shaping PLN monomer phosphorylation response at steady state.

### 2.5 Phosphorylation delay and bistability are effective noise-filters

Like the phosphorylation response delay, the emergence of bistability poses the question what the physiological advantage of such phenomenon might be Due to the small bistable range, it seems unlikely that PLN phosphorylation is a potent all-or-nothing switch as known for bistable signaling networks controlling e.g. the cell cycle or apoptosis In fact, adapting cardiac performance to various levels of demand requires the response to *β*-adrenergic stimulation to be tunable.

Altered Ca^2+^ -handling is a known cause for cardiac arrhythmias [41]. Cardiac arrhythmias such as ventricular tachycardias and fibrillation are also a hallmark of the pathogenic PLN mutation R1 del,[12, 42 -44]. We thus speculated that delayed and bistable PLN phosphorylation might play a role in preventing such arrhythmias If the phosphorylation delay is indeed a persistence sensor [37] for *β*-adrenergic stimulation, it indicates that a cardiomyocyte’s ‘decision’ to phosphorylate PLN may be a critical one We hypothesized that by controlling PLN phosphorylation, response delay and bistability are noise-filtering mechanisms to prevent random, uncoordinated *β*-adrenergic signaling and aberrant Ca^2+^ -handling.

To test this hypothesis, we performed a series of different simulations and analyses to characterize the noise handling behavior of the model in response to random fluctuations of PKA activity In the first simulations, we explored monomer phosphorylation in response to short bursts (1/ 3.3 / 0s) of maximal PKA activity (0.59 *µM*) in the full model, in the absence of either pentamers or inhibitor-1, and in the absence of both pentamers and inhibitor-1 In the full model, the first 1/3.3 /10s bursts lead to 6/13/28% monomer phosphorylation, respectively, and modestly increase thereafter (Figure 5A) In the absence of either pentamers or inhibitor-1, the response to such bursts is markedly higher, reaching 10/3.3 /46 % for 1/ 3.3/10s bursts in the absence of both pentamers and inhibitor-1 (Figure 5B-D) These simulations show that the response delay via pentamers and inhibitor-can filter out or attenuate short PKA activity bursts while still allowing high phosphorylation upon persistent PKA activity.

**Figure 5.**
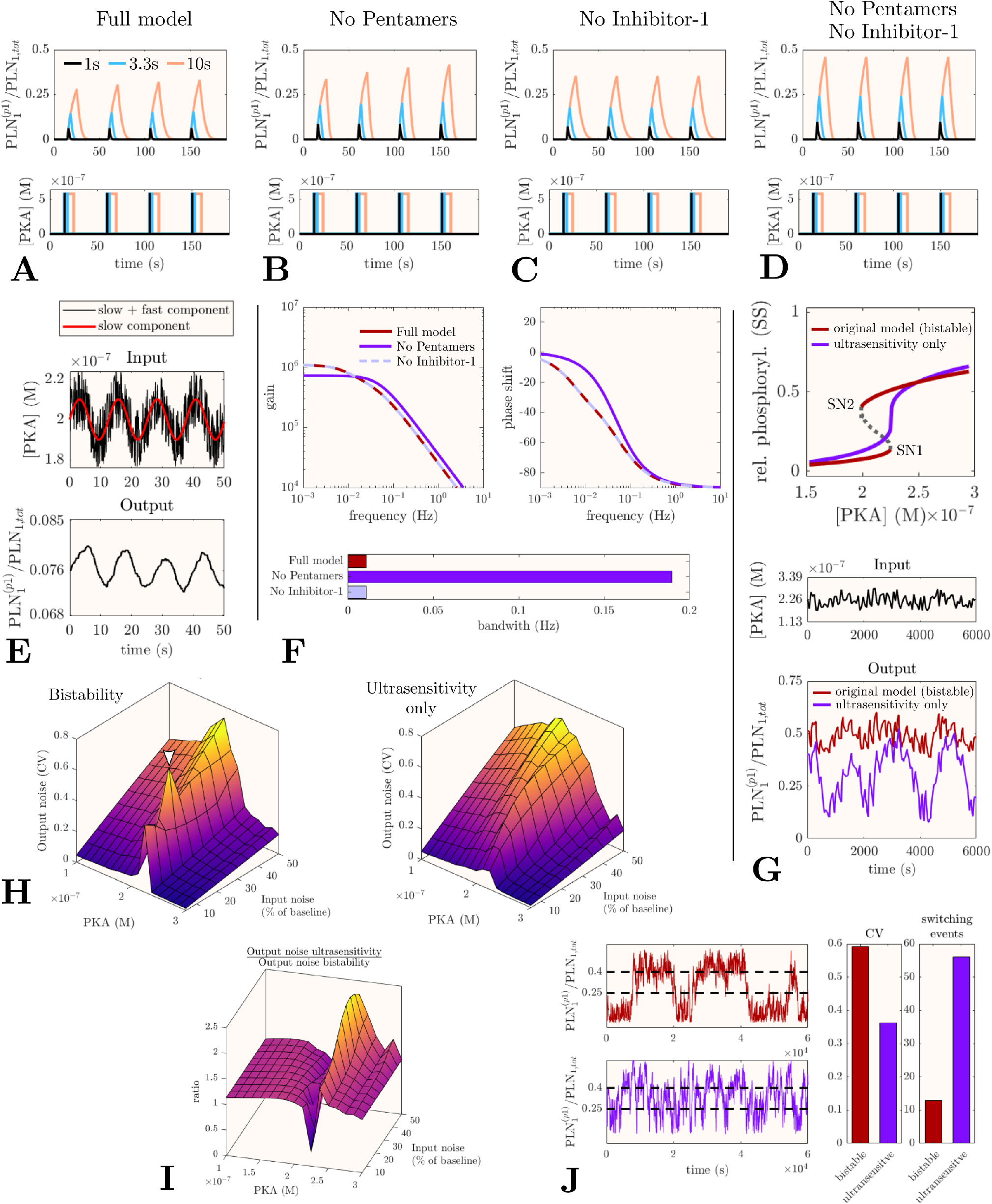
Noise-filtering by the PLN network. (A-D) Time-course simulations of PLN monomer phosphorylation in response to short (1/3.3/10s) bursts of maximal PKA activity performed with the full model (A) or in the absence of pentameric PLN and/or inhibitor-1 (B-D). To ensure equal steady-state phosphatase activity PP levels in (C, D) have been adjusted by the amount in complex with inhibitor-1 from at steady-state in the full model. (E) Demonstration of the PLN network ’s low pass filtering capacity. (F) frequency response analysis (Bode-plots) of the linearized input-output systems, (G) Comparison of PLN monomer phosphorylation (bottom) in response to a noisy PKA input fluctating with ±25% min ^-1^ around a baseline of 0.226 *µ*M (middle) for the original model and a model with similar steady-state response but without bistability (top). (H) Output noise as the coefficient of variation *σ/µ* of monomer phosphorylation for the original (bistable) and ultrasensitive model at different PKA baseline and input noise levels. (I) Output noise of the ultrasensitive model relative to the bistable model. (J) Comparison of bistable and ultrasensitive model at critical PKA concentrations and a maximum noise amplitude (0.0625*µ*M) which enables repeated switching between low/high phosphorylation (dashed lines) in both models.

Rejecting signals on short time scales while responding to persistent signals is also characteristic of low-pass filters Simulating the PLN phosphorylation response to a PKA input described by a low frequency sine wave interspersed with high-frequency random noise shows that the PLN signaling network has indeed low-pass filtering properties (Figure 5E) Such behavior can be further characterized by a frequency response analysis which allows to the determine the *bandwidth*, i e the frequency above which a system fails to response adequately Typical for low-pass filters, the gain Bode-plot of our model shows a steady decrease of the gain (roll-off) for frequencies above the bandwidth (Figure 5F) Interestingly, the bandwidth is 17 -fold higher in the absence of pentamers (0.196 Hz) compared to the full model (0.011 Hz)(Figure ,5F bar graph), confirming that pentamers contribute to the low-pass filter function of the PLN network.

To our surprise, the absence of inhibitor-1 did not increase the bandwidth (contrary to what would be expected from the demonstrated response delay) The reason for this is that the frequency response is constructed from the reached steady state Due to the high affinity of inhibitor-for PP1 ,99. 8% of inhibitor-1 at the studied steady state in the full model is already bound to PP and does not contribute to low-pass filtering anymore Unless inhibitor-can be dephosphorylated by PP2A whilst bound to PP1 (which to our knowledge has not been studied yet), its response delay would only apply if the cardiomyocyte has not been exposed to significant *β*-adrenergic stimulation for some time.

Taken together, our simulations and analyses show that the response delay by PLN pentamers and inhibitor-1 can attenuate the response to short bursts of PKA activity and that at least pentamers contribute to low-pass filtering.

Next, we explored how bistability might contribute to noise-filtering In general, bistability can make a response more robust and defined: once a system passed a threshold, it can only switch back to it’s prior state if it passes a second threshold, thus preventing uncontrolled switching [45]. We thus speculated that bistability could reduce noise by preventing repeated switching between low/high PLN phosphorylation levels To test this hypothesis, we first created a parameter set for which the model shows similar monomer phosphorylation at steady state in terms of ultrasensitivity and critical threshold but without bistability (Figure 5G, top) Next, we compared the behavior of both parametrizations in response to noisy PKA activity close to the common critical threshold Fluctuations of ± 25% with a frequency of 1 min ^-1^ have been chosen to make sure the fluctuations are not filtered out by low-pass filtering (Figure 5G, middle) As shown in Figure 5G (bottom), the relative PLN monomer phosphorylation of the bistable model (red) fluctuates with small amplitude around a stable baseline of approximately 50% phosphorylation In contrast, the non-bistable model (purple) shows dramatic fluctuations between low and high phosphorylation levels Since PLN monomer phosphorylation directly translates into SERCA activity, such fluctuations could impair coordinated Ca^2+^handling.

To investigate the output noise in a more systematic manner, we applied a common definition of signal noise as the coefficient of variation (CV)[46] By calculating ‘noise landscapes’ for both models based on 150 PKA fluctuations with a frequency of min^-1^, we visualized how the CV of monomer phosphorylation (output noise) depends both on the baseline PKA activity and amplitude of PKA fluctuations (input noise) While the output noise of bistable and non-bistable model versions is very similar for baseline [PKA] below ≈ 0.2 *µ*M, the output noise of the bistable model abruptly increases at a baseline [PKA] close to the critical threshold and abruptly decreases at higher [PKA] (Figure 5H left) The output noise of the non-bistable model follows a more continuous trend and neither shows abrupt increases close to the critical threshold, nor abrupt suppression at higher baseline [PKA] (Figure 5H, right).

The relative noise landscape shows that in most circumstances, the bistable model copes better with noisy input than the non-bistable model (Figure 5I) Since we assumed the input noise to be a linear function of the baseline [PKA], we repeated the analyses assuming a constant and a non-linear noise function and came to the same conclusion (Figure S7).

Since the bistable model seemingly performs worse in some conditions close to the critical threshold, the question arises whether this could facilitate cardiac arrhythmias in spite of a generally less noisy monomer phosphorylation To answer this question, we analysed one of the conditions in which the bistable model seemingly performs worse (white arrowhead in Figure 5H) in more detail Interestingly, we found that the increased output noise as defined by the CV typically resulted from a single ‘switching up’ event and that in the long run (1000 fluctuations), the output noise (CV) of the bistable model is actually lower than in its non-bistable counterpart (data not shown).

Motivated by this finding we wanted to know how bistable and non-bistable model versions compare at their most vulnerable point for uncontrolled switching between low/high monomer phosphorylation states We thus designed simulations in which baseline [PKA] was set to the centre between both saddle-node bifurcations for the bistable model (i e between critical thresholds SN1 and SN2 shown in Figure 5G) or directly to the single threshold in the non-bistable model. In addition, we chose a constant maximum noise amplitude for both models, high enough to surpass both thresholds in the bistable model from its baseline [PKA]. Intriguingly, we found that in spite of a higher CV, monomer phosphorylation is more defined and switches far less frequently in the presence of bistability (Figure 5J).

In summary, these simulations and analyses confirm our hypothesis that phosphorylation delay and bistability can act as molecular noise-filters in the *β*-adrenergic signaling network.

### 2.6 The R14del mutation likely impairs noise-filtering

Coordinated contraction and relaxation of the heart critically depends on the synchronicity of cardiomyocyte contraction and relaxation controlled by intra-Cellular [Ca^2+^] Since *β*-adrenergic stimulation is a major regulator of cardiac Ca^2+^ -handling, there likely need to be mechanisms in place to prevent arrhythmias triggered by heterogeneous cardiomyocyte responses. By rejecting short random stimuli (low-pass filtering) and by defining the PLN phosphorylation status more clearly (bistability), these noise-filters could help to promote synchronicity across the myocardium.

Since cardiac arrhythmias are a major issue for patients with the PLN mutation R14del [12,42-44], we wanted to know whether noise-filtering is impaired if we implement the known molecular effects of this mutation into our model The consequences reported so far include impaired phosphorylation by PKA [47] (although some studies reported PLN_*R*14*del*_ can still be partly phosphorylated *in vivo* [12,48]), mistargeting of mutant PLN to the plasma membrane [48] and destabilization of pentamers [12]. Although all R14del patients are reported to be heterozygous, explicit accounting for both wild-type and mutant PLN molecules requires a model at least three times the complexity of the current model (due to combinatorial expansion of reactions, molecules and system equations). Since this exceeds the scope of the current study as well as available data on parameters, we opted for an alternative approach and made qualitative predictions of how known molecular effects of the R14del mutation would individually influence the noise-filtering based on the analyses of the original model.

Our qualitative predictions suggested that in a heterozygous setting, mistargeting of R14del PLN, destabilization of pentamers and potential mutant/wild-type hetero-pentamers would impair both low-pass filtering and bistability (Table 1). Thus, the heterozygous R14del situation could be more permissive for short random bursts of PKA activity and lead to higher noise amplitudes, providing an attractive explanation for the susceptibility to cardiac arrhythmias. Since R14del PLN molecules are unresponsive to phosphorylation by PKA, we expect reduced amounts of wild-type pentamers to be the biggest issue for noise-filtering.

**Table 1.** Qualitative predictions for the expected influence of R14del effects on noise-filtering.

| R14del effect | low-pass filtering | bistability | based on |
| --- | --- | --- | --- |
| mistargeting of R14del PLN<br>→ lower $[\text{PLN}_{\text{tot}}]$ at SR<br>→ less pentamers | ↓ | ↓ | sensitivity analysis,<br>Fig. S3 |
| destabilisation of pentamers<br>→ less pentamers | ↓ | ↓ | sensitivity analysis,<br>Fig. S3 |
| (wildtype/R14del hetero-pentamers)<br>→ less phosphorylation sites | (↓) | (↓) | (Markevich et al. 2004) |

Although preliminary, our analysis suggests a novel therapeutic strategy: increasing the amount of wild-type pentamers could improve noise-filtering and prevent cardiac arrhythmias in patients with the R14del mutation. Potential ways of achieving this include increasing the effective concentration of PLN at the SR, small (and yet to be discovered) molecules which stabilize pentamers without interfering with regulatory enzymes or metabolically changing the lipid composition of the SR (which regulates PLN pentamerization[49]).

## 3 Discussion

In the present study, we have demonstrated that at least in our experimental conditions, the buffering effect exerted by PLN pentamers is too moderate and slow to be relevant at the time scale of acute *β*-adrenergic stimulation. We therefore developed a mathematical model of the PLN regulatory network to study the role of PLN pentamers in the context of *β*-adrenergic stimulation from a dynamical systems perspective. Having calibrated the model with own experimental data and experimental parameters from the literature, our simulations predicted delayed phosphorylation responses due to PLN pentamer competition and an inhibitor-1 FFL. Further simulations suggested that PLN phosphorylation could be ultrasensitive and bistable due to cooperative dephosphorylation of PLN pentamers.

Using several different numerical approaches we have shown that these phenomena can filter out the effect of random fluctuations in PKA activity on PLN monomer phosphorylation: while response delay and persistence sensing constitute a low-pass filter removing fast fluctuations and short stimulus spikes, bistability prevents uncontrolled high-amplitude fluctuations in PLN phosphorylation at critical PKA activity, thereby promoting a well defined PLN phosphorylation status. Importantly, these noise-filters are complementary (Figure 6) and depend largely on PLN pentamers. To our knowledge, this is the first time that a clearly defined physiological advantage of PLN pentamers has been demonstrated.

**Figure 6.**
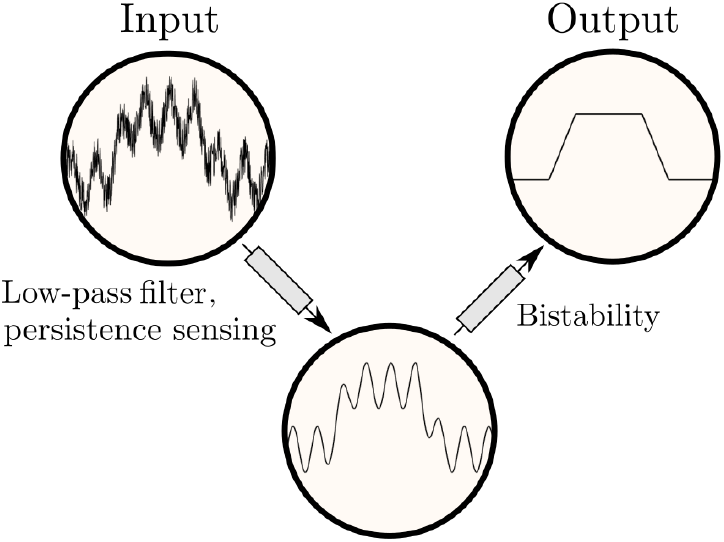
Illustration of complementary noise-filtering processes in the PLN signaling network.

While we have provided an optimal design for experimentally testing the FFL functionality e.g. with cell biological approaches, we have confirmed the delay due to pentamer competition, providing experimental evidence for one of the main mechanisms underlying the predicted noise-filtering. Similar monomer phosphorylation delays due to pentamers have been observed in transfected HEK293 cells, suggesting the mechanism can operate in living cells[25].

Using the same experimental approach, we further confirmed ultrasensitive PLN phosphorylation in transfected EK cells This demonstrates that PLN phosphorylation meets an important prere uisite for bistability Re-analysing previous experimental data by fitting it to a Hill-equation showed that this ultra-sensitivity depends on the presence of pentamers (Figure S8), likely due to the (pseudo-)multisite nature of pentamers [38] Although the simulated dose-responses appear steeper than the experimental ones, we note that the experimental Hill-exponents may be underestimates due to limited dose resolution and averaging out of the response across cell populations However, lower ultrasensitivity, too, is perfectly consistent with the presented findings (Figure S9) In an independent study we show further that the ability of pentamers to shape the response curve of PLN phosphorylation to *β*-adrenergic stimulation translates into increased dynamic range and sensitivity of cardiac relaxation and is necessary to cope with increased cardiac pressure [50].

Our results also provide an explanation for the frequent emergence of cardiac arrhythmias in patients with the R14 del mutation Although preliminary, a first analysis of the molecular consequences of this mutation indeed points towards impaired noise-filtering due to reduced amounts of wild-type pentamers Since the PLN_*R*14*del*_ cardiomyopathy does not respond to conventional heart failure therapy [51], we propose to explore ways to increase the amount of wild-type pentamers in preclinical R14del models as a novel therapeutic approach A first way to test this concept could be to harness increased pentamerization of the artificial I45A mutant and to determine whether heterozygous R14del/I45 A mice or iPSC-cardiomyocytes show a lower arrhythmogenic tendency than a heterozygous R14 del/wild-type model.

A strength of our model is that it casts new perspectives on multiple and hitherto puzzling phenomena by associating them with a common physiological function: noise-filtering Apart from PLN pentamers, neither their cooperative dephosphorylation [23,35], nor the dynamic equilibrium (increased pentamerization upon phosphorylation [30,31]) were previously known to have clearly defined physiological functions While pentamers and their cooperative dephosphorylation are necessary in our model for bistability to occur, the dynamic equilibrium makes bistability more robust, potentially by inducing ‘hidden’ feedback loops [52] supporting the emergence of bistability (Figure S10) Menzel *et al* recently found that recruitment of 14-3-3 proteins to phosphorylated PLN pentamers generates a similar memory of PLN phosphorylation *in vivo* (by slowing down dephosphorylation) and is impaired by R14del, too [53] This constitutes a double-negative feedback which may further improve bistability or noise-filtering.

Our model also casts a new perspective on the role of inhibitor-1, which is often described as an amplifier for PKA phosphorylation[54,55] Although this is technically not false, the same level of PLN phosphorylation response could in principle be achieved by simpler means such as reduced SR targeting of PP1[56] Thus, the delayed response dynamics of the inhibitor-1 FFL and the noise-filtering capacity demonstrated in our simulations may be equally important as the influence on steady-state phosphorylation levels.

### 3.1 Further experimental and clinical evidence

Noise-filtering in the *β*-adrenergic signaling pathway may only be relevant if the network indeed experiences significant fluctuations which can be pro-arrhythmogenic under some conditions While it is known that there is significant electro-physiological variability among individual cardiomyocytes which can be pro-arrhythmogenic under conditions of reduced cell-cell coupling [57], a systematic experimental characterization of the noise at multiple nodes of the *β*-adrenergic signaling network is, to our knowledge, still lacking However, studies from the 80s indicate significant fluctuations at baseline at least in cate-cholamines [58,59] and aberrant calcium-handling or *β*-adrenergic signaling are well known to be able to trigger cardiac arrhythmias 4[1], making noisy signaling in these pathways a hypothesis worthwhile to study further.

In agreement with our results, many proteins of the PLN regulatory network are in fact associated with cardiac arrhythmias by both experimental and clinical data The natural mutation R9H[13] has recently been shown to cause ventricular arrhythmias in dogs[14], indicating that PLN mutations other than R14 del can be arrhythmogenic Additional evidence that PLN pentamers contribute to noise-filtering comes from a mouse model of the natural obscurin variant R4344Q[60] Mice carrying this variant developed spontaneous ventricular arrhythmias which authors attributed to increased SERCA levels and ≈15% less pentamers Although the pathogenicity of this variant is likely restricted to mice[61], the findings support the idea that pentamers can attenuate cardiac arrhythmias A similar mechanism may contribute to the pro-arrhythmic effect of thyroid hormones which increase SERCA and decrease PLN expression (thus decreasing pentamerization) [62] Apart from PLN, both PP1 and inhibitor-1 have been shown to be involved in the emergence of arrhythmias [63] Reducing the concentration of PP1 at the SR by ablating its targeting subunit PPP1R3A has been shown to lead to atrial fibrillation [56], consistent with a smaller bistable region expected from reducing [PP1] in our model Interestingly, a mouse model of the human inhibitor-1 variant G109E (showing reduced binding to PP1) and mice expressing a constitutively active version of inhibitor-1 developed severe cardiac arrhythmias in response to *β*-adrenergic stimulation 64,65] Both mutations interfere with the inhibitor-1 FFL and could make PLN phosphorylation more susceptible to noise according to our model Contrary to these findings, complete inhibitor-1 ablation has been shown to protect against catecholamine-induced arrhythmias[66] which led to a debate over whether inhibitor-1 is pro- or anti-arrhythmogenic[55,67] Settling this debate may require a nuanced answer distinguishing between pro- and anti-arrhythmogenic effects (e g pro-arrhythmogenic PKC phosphorylation sites vs noise-filtering) Moreover, the complete loss of inhibitor-1 may be compensated by up-regulating FFLs involving e g Hsp 20 [8].

An intriguing line of evidence that noise-filtering may also be up-regulated in response to arrhythmic heart activity came from a recent study on arrhythmogenic cardiomyopathy (AC) patients In AC patients without PLN mutation, PLN protein expression was shown to be up-regulated more than twofold, which the authors hypothesized to be a yet to be elucidated compensatory mechanism [68] As higher PLN concentration leads to increased pentamerization due to mass action, both noise-filtering functions predicted by our model would be enhanced, providing an attractive explanation for this observation.

Taken together, these studies show that perturbing components which contribute to noise-filtering in our model can lead to cardiac arrhythmias, whereas enhancing their functionality may protect against other pro-arrhythmogenic factors.

### 3.2 Limitations and concl sion

Like every modeling study, we had to rely on simplifying assumptions at several points during model development For example, our model only accounts for a subset of interactions with PLN which we deemed most relevant for our purpose; it assumes that the modeled processes are described well enough by ordinary differential equations even though much of the biochemistry takes place on the two dimensional SR surface; parameters and species concentrations of our model come from different sources (e g fitted to own experimental data and directly measured or fitted parameters from the literature) Furthermore, we have considered noise only in terms of fluctuating PKA activity (representing the ‘input’ of our model) although extrinsic and intrinsic noise sources affect all molecular processes whose full exploration re uires stochastic simulations[46,69] Due to its medical relevance, the perhaps most important limitation to highlight is that our analysis of the R14del mutation is preliminary Since this mutation has been shown to also alter other interactions not currently represented in our model, the model does not yet capture other pathophysiological processes such as (potentially pro-arrhythmogenic) cardiac remodeling [70].

While we have partly addressed some limitations e g by performing complementary analyses or systematic explorations of the parameter space, others will need to be addressed in future experimental and theoretical investigations Despite these limitations, we believe our model offers a novel and exciting perspective on the physiological role of PLN pentamers that will prove to be a useful starting point for further investigations.

## 4 Methods

Detailed descriptions of the model as well as experimental and computational procedures applied in this work can be found in the online supplementary material.

The model code will be accessible in the Bio odels database upon publication at: https://www.ebi.ac.uk/biomodels/MODEL.2011110001.

## Acknowledgements

DK is funded by a PhD studentship from the British Heart Foundation (grant [FS /17/ 65/33481]) we would like to thank Dr. Thomas Kampourakis and Dr. Martin Rees for helpful criticism on earlier versions of the manuscript.

## Author contributions

DK designed the study formulated the mathematical model and performed simulations AA,DK and FF performed experiments and analysed the data DK wrote the manuscript JPS and MG provided important feedback and revised the manuscript.

