## Supplementary Material for "Molecular Noise-Filtering in the *β*-adrenergic Signaling Network by Phospholamban Pentamers" (from another version of this preprint: version 1, 2020-12-30)

---

**4 Appendix: simulation protocols** **32**

---

### 1 Supplementary figures

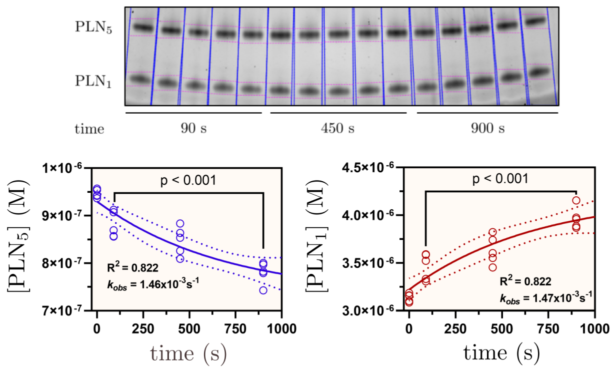

**Figure S1**
PLN dissociation time course. Top: pre-equilibrated PLN at a concentration of ≈ 1 mg/ml was diluted 20-fold and dissociation was allowed to proceed for 90 to 900 s (each experiment was started at different time points to ensure a simultaneous endpoint). Bottom: quantification shows a very slow, but statistically significant ($p < 0.001$) pentameter dissociation and monomer accumulation. Concentrations at $t = 0$ s were calculated from the equilibrium concentration of PLN at 1 mg/ml by division through 20, i.e. assuming no dissociation directly after dilution at $t = 0$ s.

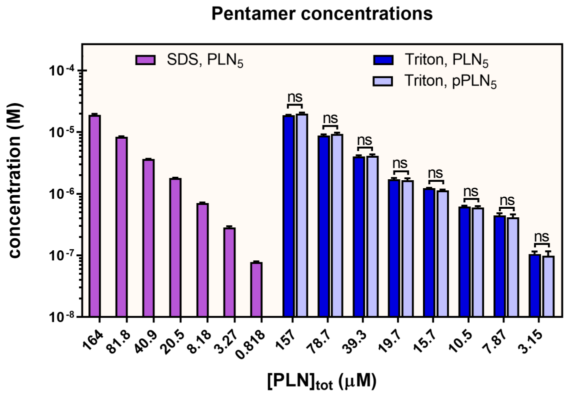

**Figure S2**
PLN pentamer concentrations at different total PLN concentrations show no significant differences between unphosphorylated and phosphorylated PLN in TBB (same data as in Figure 1 from the main text).

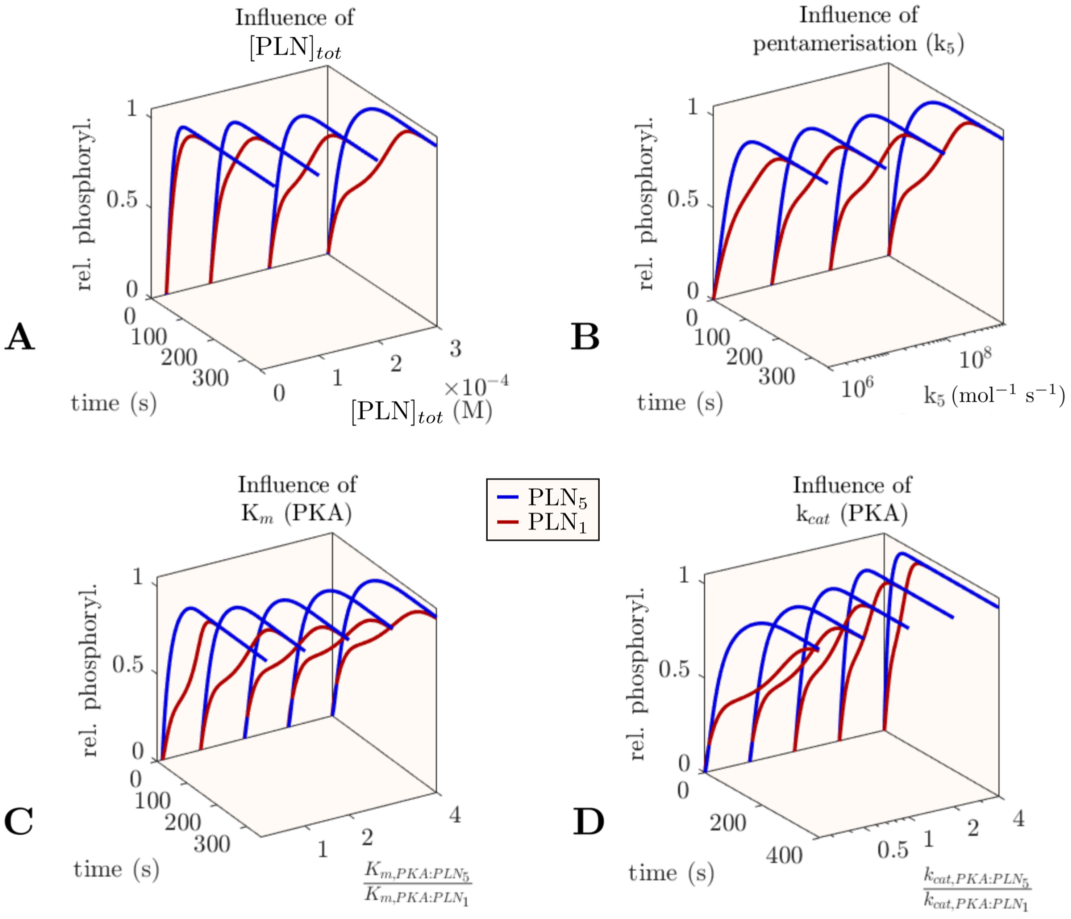

**Figure S3**
Parameter dependency of the observed monomer phosphorylation delay. (A,B) Higher [PLN]$_{tot}$ or pentamerization contribute to the monomer phosphorylation delay, likely by increasing the competitive effect of pentamers. (C,D) Monomer phosphorylation delay is more pronounced if pentamers are a worse substrate for PKA than monomers (i.e. either lower substrate affinity $K_m$ or lower turnover number $k_{cat}$). Although surprising at first glance, this becomes plausible considering that the competitive effect must abate once all pentamers have been phosphorylated.

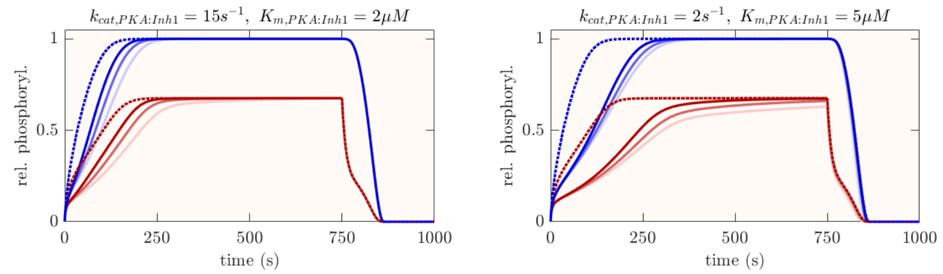

**Figure S4**

Response delays by inhibitor-1 FFL at slower phosphorylation rates of inhibitor-1 by PKA.

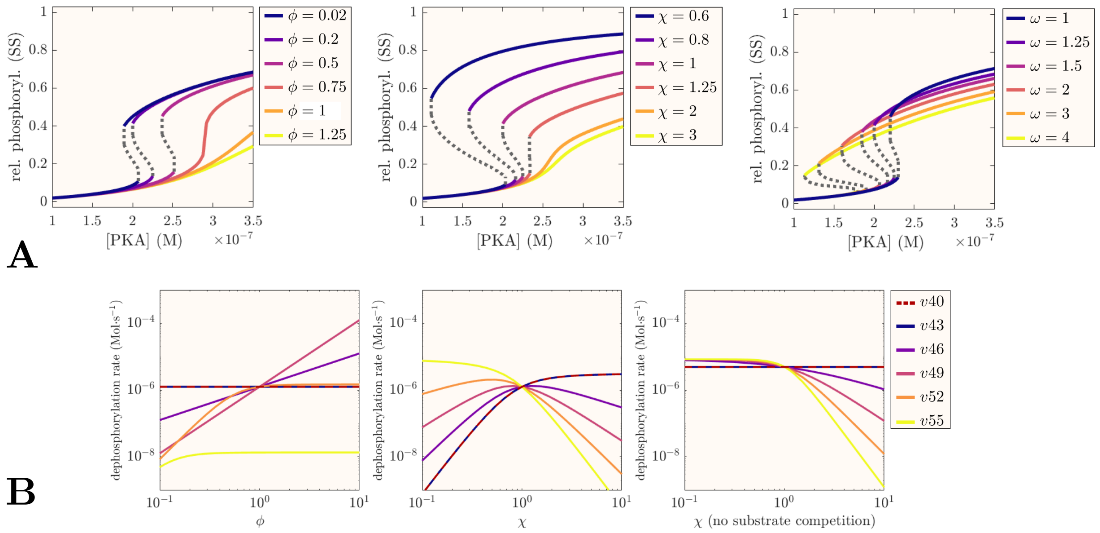

**Figure S5**

(A) Bifurcation plots of relative PLN monomer phosphorylation for different strengths and types of PP1 cooperativity ($\phi, \chi$) and for the dynamic equilibrium of PLN ($\omega$). Parameters, which were not varied, are equal to their default value (Table S3). (B) PP1 dependent dephosphorylation rates for monomeric and pentameric PLN as a function of parameters $\phi$ and $\chi$.

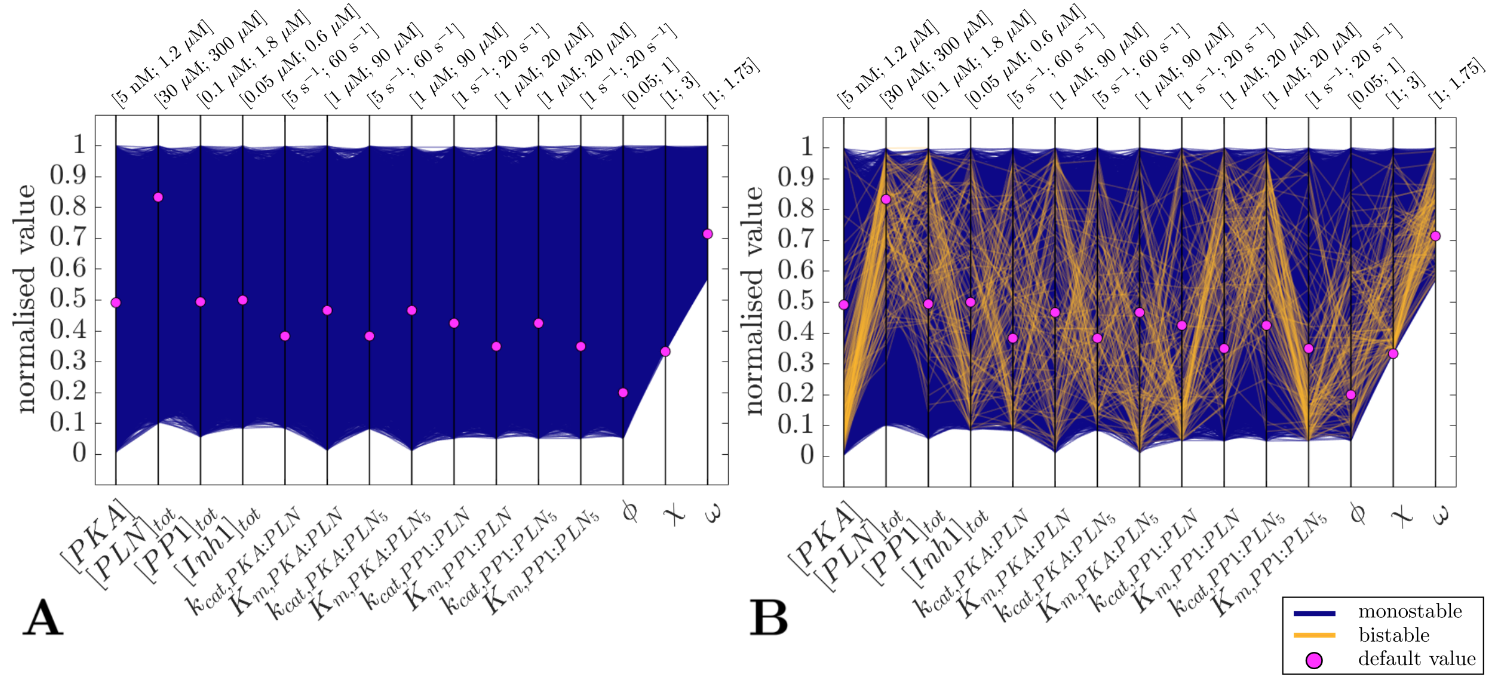

**Figure S6**
(A) No bistability could be observed in the absence of pentamers regardless of the values for other kinetic parameters or protein concentrations. (B) Implementing k-type cooperativity (parameter $\chi$) reduces the probability for a parameter set to be bistable as only 1.1% of tested parameter sets showed bistability.

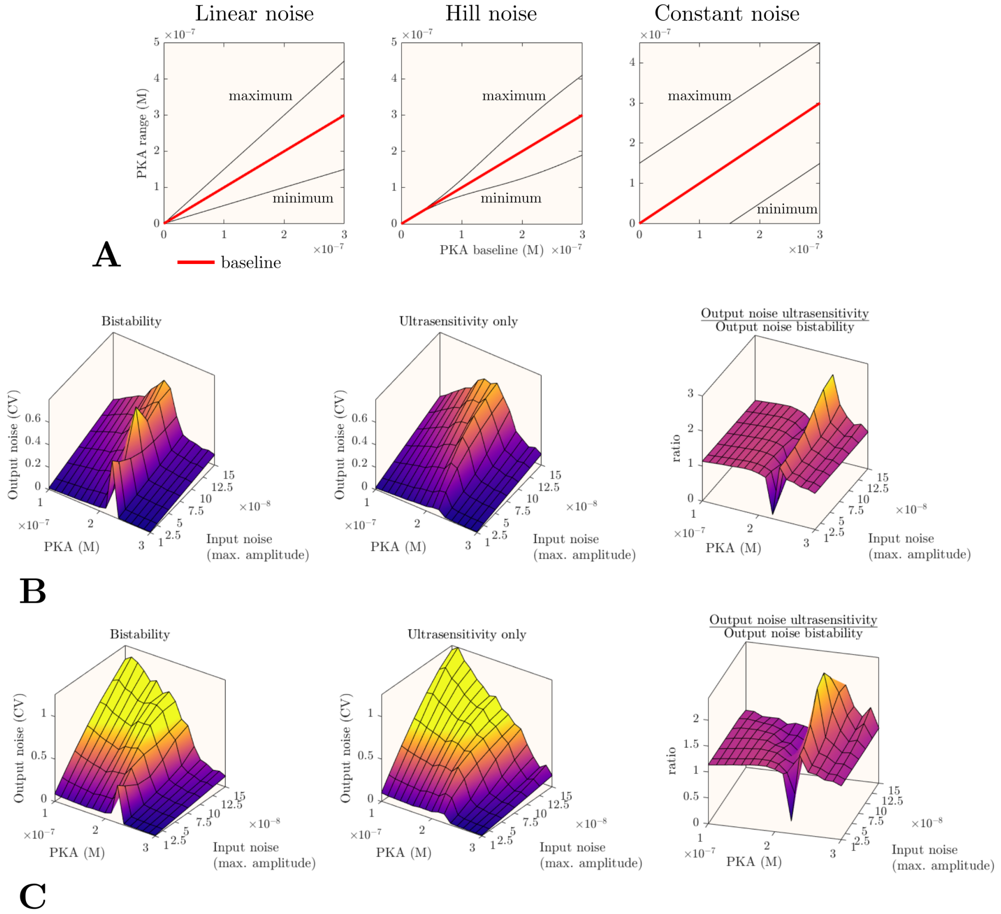

**Figure S7**

Alternative noise functions. (A) Left: linear noise underlying the noise landscapes in the main text. Middle: non-linear noise amplitude as a Hill-function of the baseline [PKA] with $K = 2 \times 10^{-7}$ M and $n_H = 2.5$. Right: constant noise amplitude. (B) Absolute and relative noise landscapes with Hill noise. (C) Absolute and relative noise landscapes with constant noise amplitude.

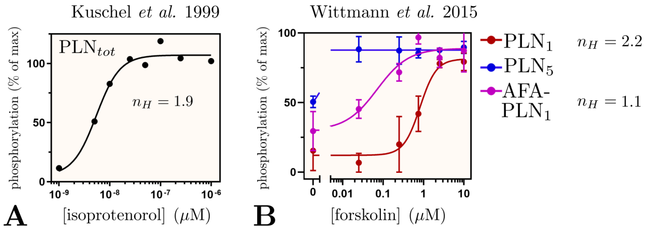

**Figure S8**

Dose-response data on PLN phosphorylation from the literature was fitted to the Hill-equation with offset. (A) Phosphorylation of total PLN (pentamers were dissociated by boiling) in rat hearts [1]. (B) Differential phosphorylation of PLN monomers and pentamers in transfected and forskolin stimulated HEK293 cells [2]. Note that high Hill-exponents depend on the presence of pentamers. Although fitted Hill-exponents indicate some degree of ultrasensitivity ($n_H \approx 2$), the true response on single cell level could be blunted due to averaging [3, 4].

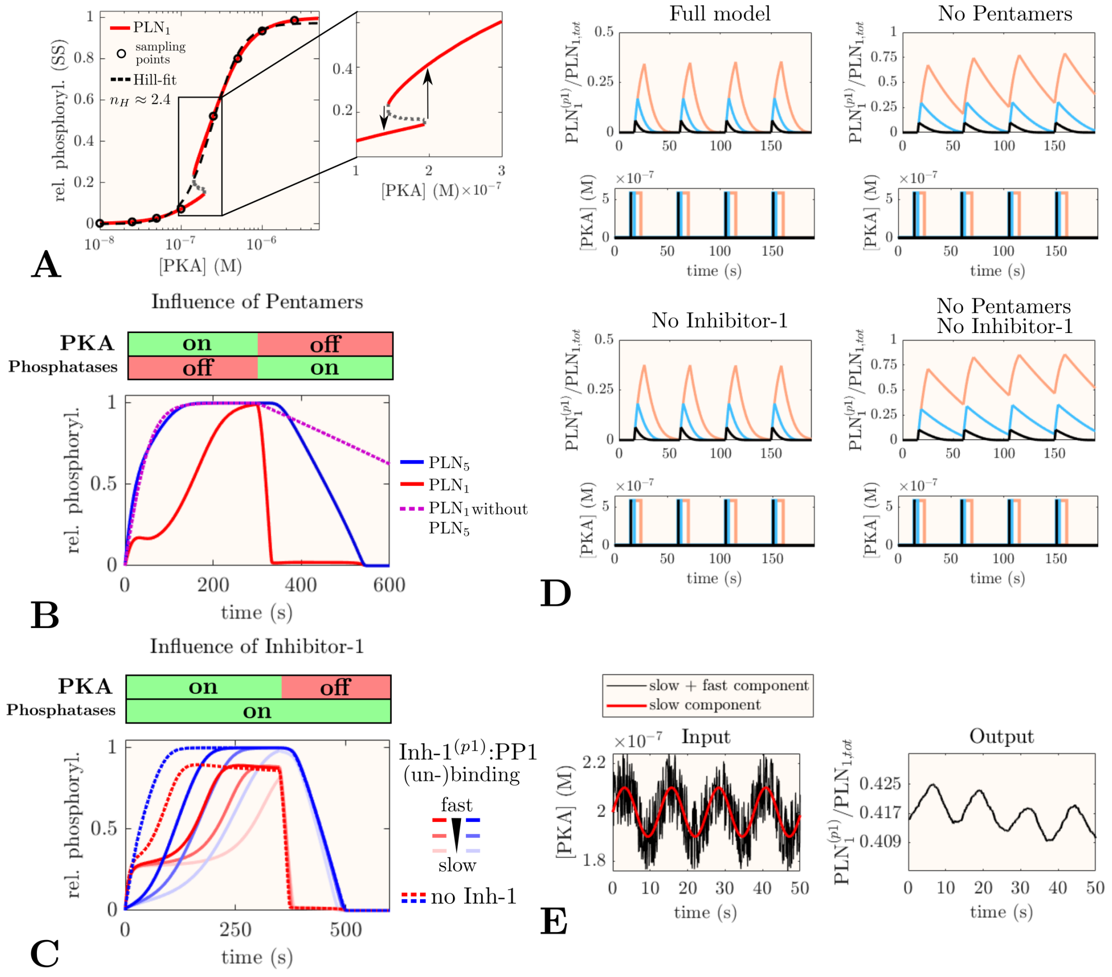

**Figure S9**

Alternative parameter set with lower ultrasensitivity. (A) Sampling the bifurcation diagram at 8 different points leads to a fitted dose-response curve with a Hill-exponent of $n_H = 2.4$ similar to experimental dose-response data. The parameter set still allows for bistability (with even higher bistable range than with the default parameters) and thus is able to filter out small fluctuations close to the critical threshold. (B,C) The alternative parameter set leads to stronger phosphorylation delay capacities than the default parameter set. Consequently, short PKA activity bursts are filtered out (D) and the PLN network can still act as a low-pass filter (E).

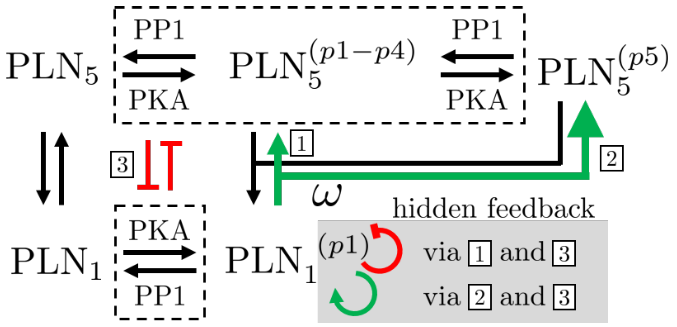

**Figure S10**
Since PLN monomers and pentamers compete with each other to be (de-)phosphorylated by PKA and PP1 (3), the dynamic equilibrium could potentially induce hidden feedback loops. First, oligomerization of phosphorylated monomers is in itself a negative auto-loop reducing the amount phosphorylated monomers. If incompletely phosphorylated pentamers are formed (1), monomer phosphorylation could be further decreased due to more pentamers competing with monomers for PKA (3). However, pentamers also compete with monomers for PP1 and could decrease monomer dephosphorylation - particularly if completely phosphorylated pentamers are formed (2). Which of these loops dominates likely depends on the relative reaction velocities among phosphorylation/dephosphorylation steps in a given situation. Hidden feedback loops were proposed to support the emergence of bistability [5].

### 2 Development of the phospholamban model

In order to develop a mathematical model of the PLN regulatory network, we first needed to find a mathematical description of how PLN monomers assemble into pentamers. Therefore, we considered several mass action kinetics based possibilities.

#### 2.1 A mass action kinetics model of PLN pentamer assembly

The assembly of PLN monomers into pentamers likely follows either of three possible pathways depicted in the scheme on the left side of Figure S11. While model 1 considers all reaction routes possible, model 2 and model 3 assume pentamer assembly to follow a monomer→dimer→tetramer →pentamer or monomer→dimer→trimer→pentamer pathway, respectively. Dimers, trimers and tetramers have been reported *in vitro* (see e.g. [6]), but monomers and pentamers are typically the predominantly observed molecular species (including the present study). While this implies that PLN oligomers with $<5$ protomers are usually low abundant and transient species, it is difficult to decide between any of the three pathways on *a priori* grounds. We thus formulated mass-action kinetics models for each possibility (Figure S11).

While the schemes for model variant 2 and 3 are linear reaction routes, model 1 contains two cycles. For thermodynamic reasons, model 1 must thus obey the following relations between equilibrium constants:

$$K_{2,3}K_{3,5} = K_{2,4}K_{4,5} \quad (1)$$

$$K_{1,2}K_{2,4} = K_{2,3}K_{3,4} \quad (2)$$

$$K_{1,2}K_{3,5} = K_{3,4}K_{4,5} \quad (3)$$

Replacing equilibrium constants with the rate constants from Figure S11 yields:

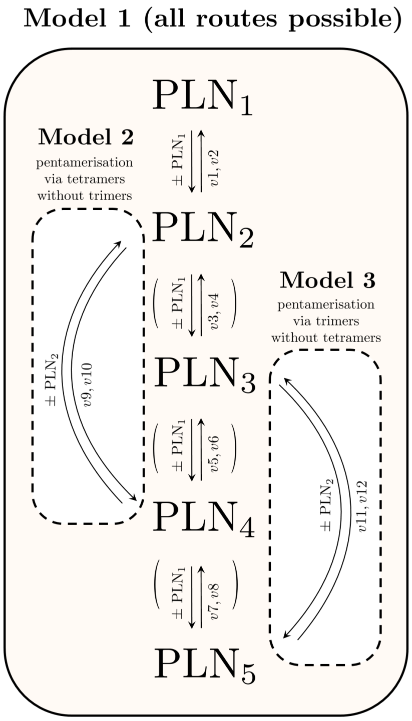

**reaction rates**

$v1 = k_1[PLN]^2$
$v2 = k_2[PLN_2]$
$v3 = k_3[PLN_2][PLN]$
$v4 = k_4[PLN_3]$
$v5 = k_5[PLN_3][PLN]$
$v6 = k_6[PLN_4]$
$v7 = k_7[PLN_4][PLN]$
$v8 = k_8[PLN_5]$
$v9 = k_9[PLN_2]^2$
$v10 = k_{10}[PLN_4]$
$v11 = k_{11}[PLN_3][PLN_2]$
$v12 = k_{12}[PLN_5]$

**equilibrium constants**

$K_{1,1} = \frac{k2}{k1} = \frac{[PLN_2]}{[PLN][PLN]}$

$K_{1,2} = \frac{k4}{k3} = \frac{[PLN_3]}{[PLN][PLN_2]}$

$K_{1,3} = \frac{k6}{k5} = \frac{[PLN_4]}{[PLN][PLN_3]}$

$K_{1,4} = \frac{k8}{k7} = \frac{[PLN_5]}{[PLN][PLN_4]}$

$K_{2,2} = \frac{k10}{k9} = \frac{[PLN_4]}{[PLN_2][PLN_2]}$

$K_{2,3} = \frac{k12}{k11} = \frac{[PLN_5]}{[PLN_2][PLN_3]}$

**ODEs**

$$\frac{d}{dt}[PLN_1](t) = 2 \cdot v2 + v4 + v6 + v8 - 2 \cdot v1 - v3 - v5 - v7$$

$$\frac{d}{dt}[PLN_2](t) = v1 + v4 + 2 \cdot v10 + v12 - v2 - v3 - 2 \cdot v9 - v11$$

$$\frac{d}{dt}[PLN_3](t) = v3 + v6 + v12 - v4 - v5 - v11$$

$$\frac{d}{dt}[PLN_4](t) = v5 + v8 + v9 - v6 - v7 - v10$$

$$\frac{d}{dt}[PLN_5](t) = v7 + v11 - v8 - v12$$

$k_3 = k_4 = k_5 = k_6 = k_{11} = k_{12} = 0$ for **Model 2**
$k_5 = k_6 = k_7 = k_8 = k_9 = k_{10} = 0$ for **Model 3**

**Figure S11**
Mass action kinetics based models of PLN pentamerization. The index denotes different oligomer size, $PLN_1$ for monomers, $PLN_2$ for dimers and so forth. Model 1 considers all reaction routes possible, whereas model 2 assumes a monomer→dimer→tetramer→pentamer and model 3 a monomer→dimer→trimer→pentamer pathway.

$$\frac{k_3 k_{11}}{k_4 k_{12}} = \frac{k_7 k_9}{k_8 k_{10}} \tag{4}$$

$$\frac{k_1 k_9}{k_2 k_{10}} = \frac{k_3 k_5}{k_4 k_6} \tag{5}$$

$$\frac{k_1 k_{11}}{k_2 k_{12}} = \frac{k_5 k_7}{k_6 k_8} \tag{6}$$

For parameter estimation, $k_7$ and $k_5$ were assigned by solving equations (4) and (5), respectively. Equation (6) was used as an additional constraint on the parameter space.

After setting up the constraints for model 1, we used the data from our dilution and dissociation time course experiments for calibration of all three models. We found that all three models can reproduce the dissociation time course sufficiently (Figure S12A). Model 1 and model 2 show generally good agreement with the data from dilution experiments apart from small deviations from the measured pentamer concentrations at the lower range of total concentrations (Figure S12B). Model 3, on the other hand, shows a systematic deviation from most experimentally measured pentamer concentrations. Simulated steady state concentrations of monomers and pentamers are fairly similar for all three model variants (Figure S12C), indicating that the shortcoming of model 3 is a mismatch at the time scale of hours. Also note that at steady state (which in this case is identical to the equilibrium of the reaction) and at $[PLN]_{tot} > 100$ µM, there is effective monomer buffering (Figure S12C, red curves).

Moreover, model 3 predicts high trimer concentrations at steady state which should have been clearly visible in our experimental conditions. Taken together, this indicates that model 3, solely relying on the monomer→dimer→trimer→pentamer pathway, cannot account for our experimental data. In contrast, although the available data are not sufficient to identify individual rate constants (Figure S13), we conclude that both model 1 and 2 can faithfully reproduce most of our experimental data. To choose between models (using their best fit parameter sets), we applied the Akaike Information Criterion (AIC) which ranked model 2 as best performing (AIC scores: model 1 = -735.2, model 2 = -741.2, model 3 = -735.9). We, therefore, used model 2 as the basis for the extended model in the remainder of this study.

Before expanding the model, we used our quantitative description of the pentamerization reaction to re-evaluate some of the numbers published on what the effective concentration of PLN in the SR membrane might be and what fraction of SERCA will be occupied at this concentration (Figure S14). Although PLN is already considered to be an abundant molecule with estimates for the effective concentration ranging from 38 to 106 $\mu$M (cf. Table S3), our analysis indicates that the effective concentration could be as high as 250 $\mu M$.

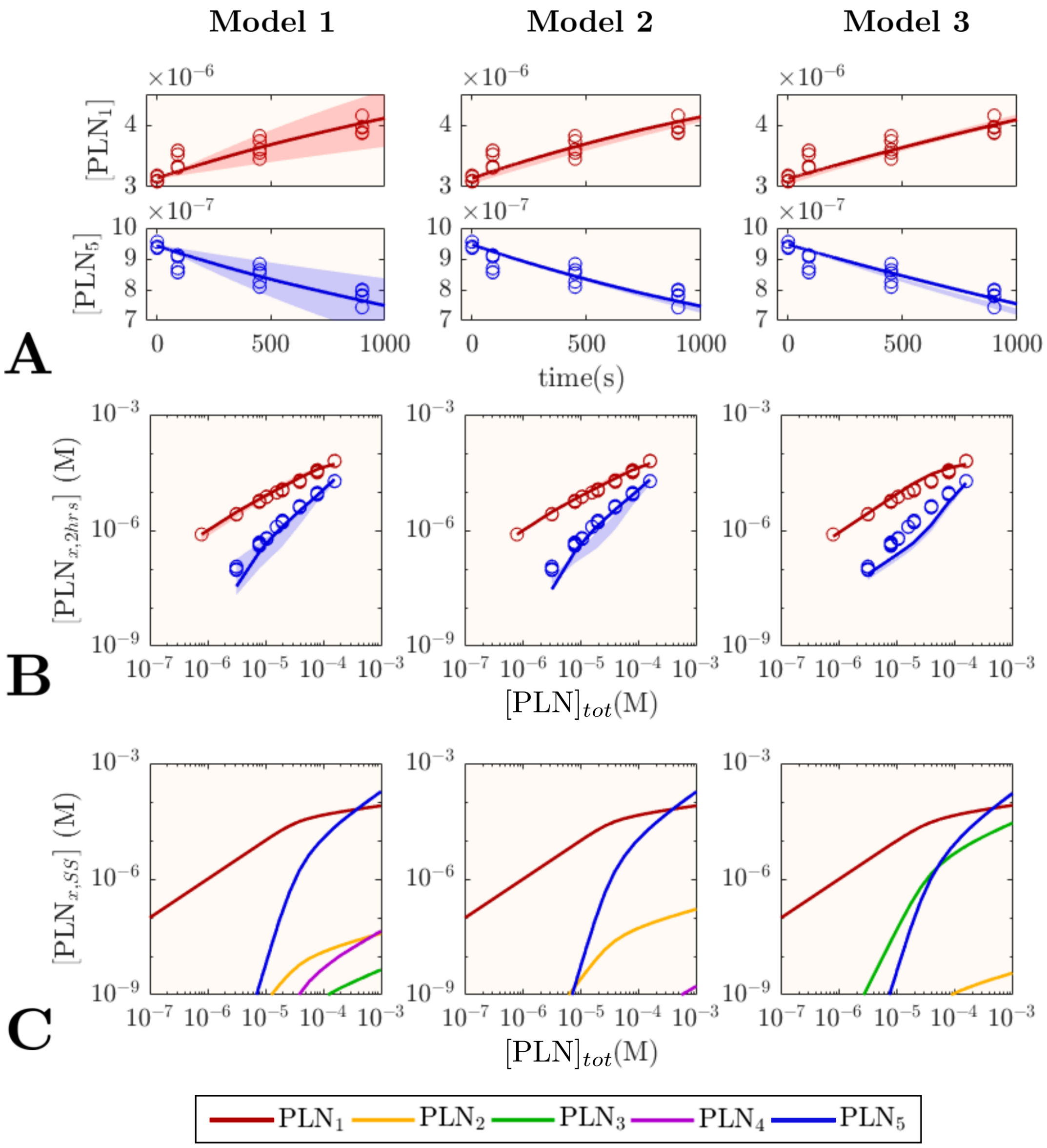

**Figure S12**
Model fits to experimental dissociation time-course (A) and dilution (B) data. Continuous lines represent results from the best fit parameter sets, circles are experimental data points, shaded areas represent 95% confidence intervals from the ten best parameter sets. (C) Simulated equilibrium concentrations of monomeric and oligomeric PLN at different total PLN concentrations using the best fit parameter set for each model.

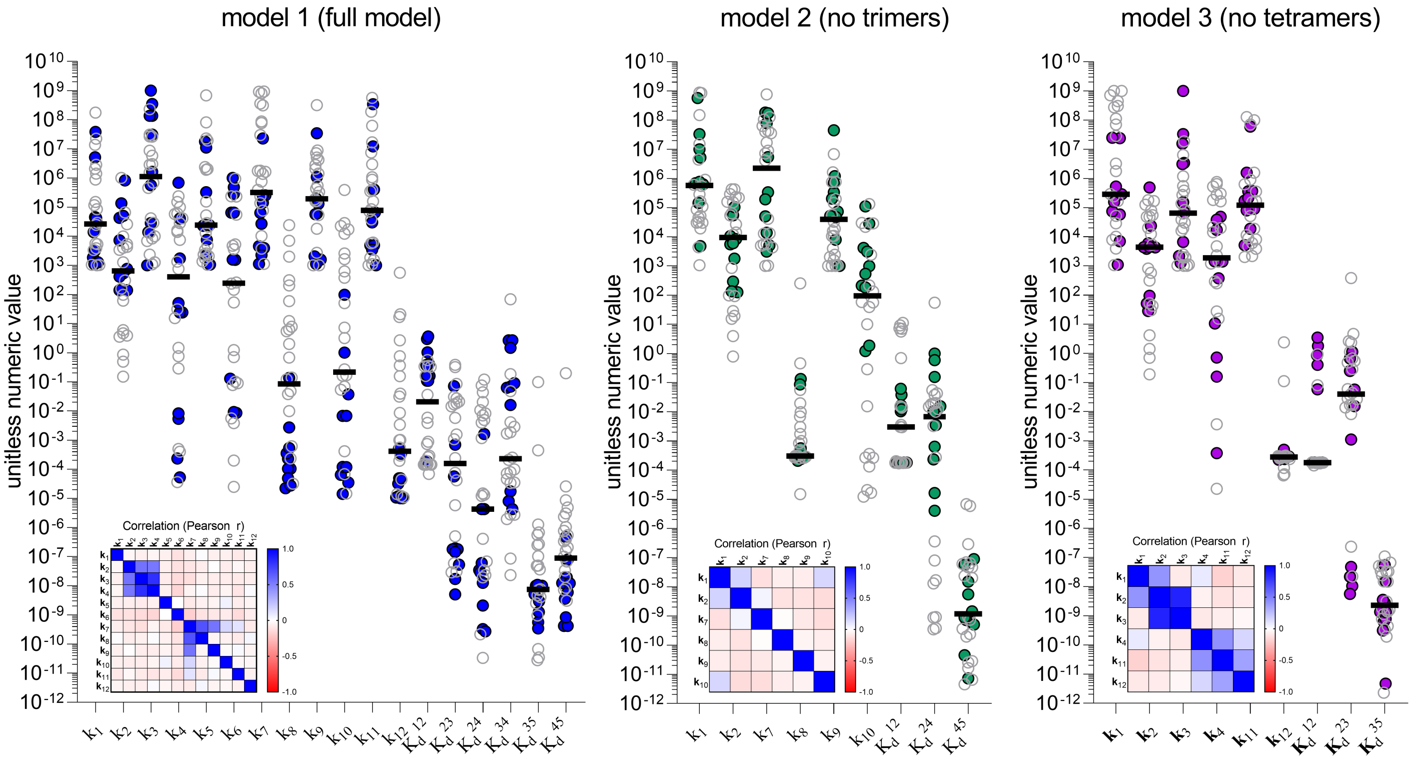

**Figure S13**
Distribution of fitted parameter values of 30 independent parameter estimation runs for model 1-3. Parameter values associated with the ten best fits are highlighted in colour. Inset shows correlation matrix of the estimated rate constants.

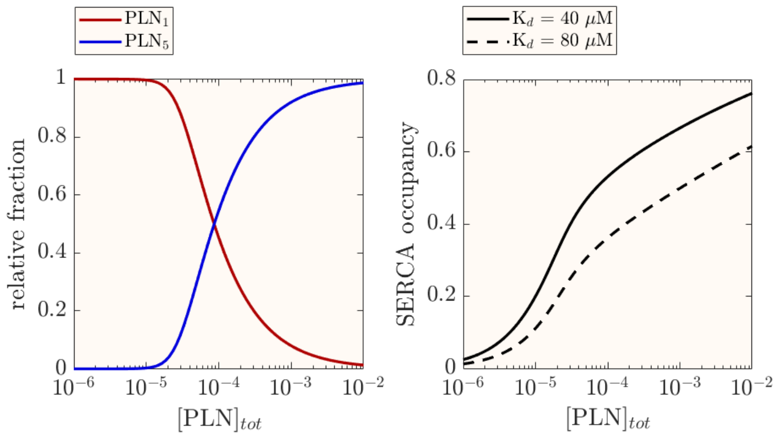

**Figure S14**
Left: relative fraction of monomers and pentamers as a function of total PLN concentration based on the best fit parameter set for pentamerization model 2. Right: SERCA occupancy by PLN assuming only monomers bind SERCA and that [SERCA] ≪ [PLN] so that SERCA binding does not influence pentamerization.

*Relevance for assessing the physiological concentration of PLN at the SR membrane*: in SDS-PAGE analyses of cell/tissue homogenates, 75-90% of PLN is pentameric [7, 8]. If oligomerization parameters of PLN in the SR membrane are similar to those in detergent micelles, this would require PLN concentrations of 200 to 600 $\mu M$ *in vivo*. Although this appears to be very high, PLN is known to be very abundant (cf. Table S1) and the required number of molecules to reach high *effective concentrations* is likely lower on a 2-dimensional surface such as the SR membrane than in solution [9]. Moreover, a PLN concentration of ≈ 250 $\mu M$ would result in a fractional SERCA occupation of about 40-60% according to published $K_d$ values. This fits well to the observation that about 40% of SERCA activity is functionally regulated by PLN [10]. At a total PLN concentration of 250 $\mu M$, the concentration of monomers is ≈ 58 $\mu M$, the concentration of pentamers ≈ 38 $\mu M$, i.e. about 76% of PLN molecules would be in a pentameric complex.

#### 2.2 Full phospholamban model in the context of $\beta$-adrenergic stimulation

As outlined in the main text, we extended the mass action kinetics model of pentamerization by accounting for PLN phosphorylation at serine 16. Phosphorylation of phospholamban leads to a combinatorial expansion of oligomeric phospho-isoforms for which we denote the number phosphorylated subunits by a superscript $(pX)$, e.g. $\mathrm{PLN}_1^{(p0)}$ for unphosphorylated monomers or $\mathrm{PLN}_5^{(p3)}$ for pentamers with 3 phosphorylated subunits etc. We further included reactions and molecules with a well established role in regulating PLN phosphorylation at serine 16, i.e. enzymes PKA, PP1, PP2A and inhibitor-1. Figure S15 shows the complete reaction scheme of our model. Before substituting rate identifiers with their rate laws, the model equations are as follows:

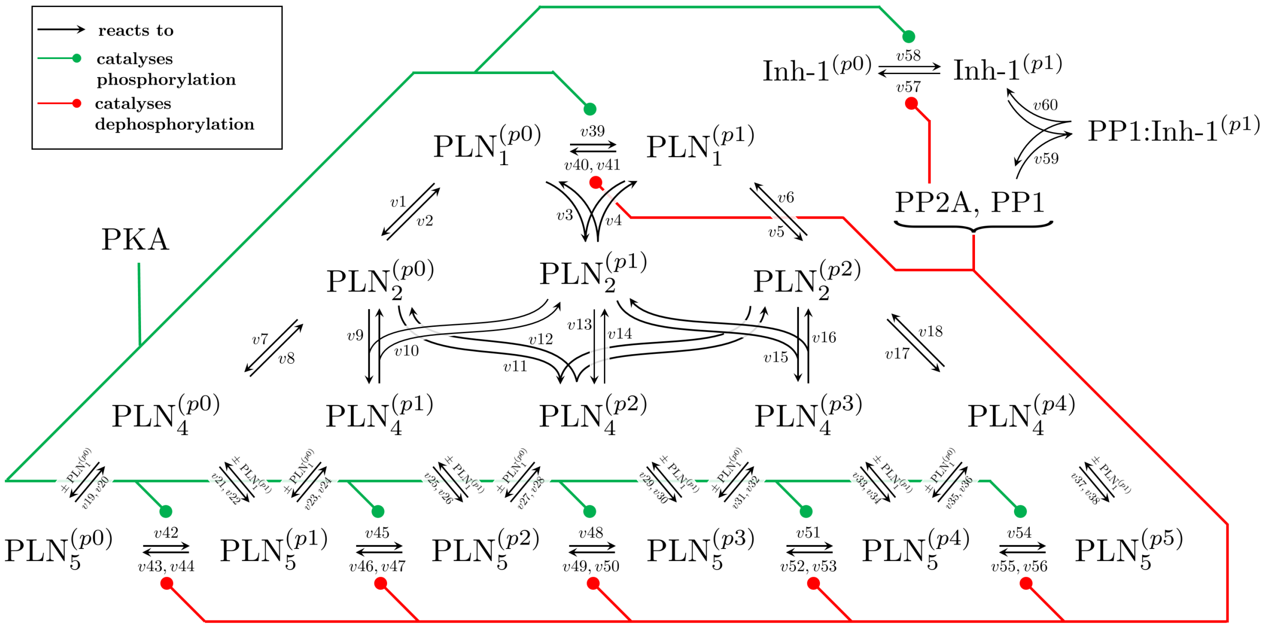

**Figure S15**

Complete reaction scheme of the phospholamban model.

#### Model equations

$$
\begin{aligned}
\frac{d}{dt}[PLN_1^{(p0)}](t) &= 2v_2 + v_4 + v_{20} + v_{24} + v_{28} + v_{32} + v_{36} + v_{40} + v_{41} \\
&\quad - 2v_1 - v_3 - v_{19} - v_{23} - v_{27} - v_{31} - v_{35} - v_{39} \\
\frac{d}{dt}[PLN_2^{(p0)}](t) &= v_1 + 2v_8 + v_{10} + v_{12} - v_2 - 2v_7 - v_9 - v_{11} \\
\frac{d}{dt}[PLN_2^{(p1)}](t) &= v_3 + v_{10} + 2v_{14} + v_{16} - v_4 - v_9 - 2v_{13} - v_{15} \\
\frac{d}{dt}[PLN_2^{(p2)}](t) &= v_5 + v_{12} + v_{16} + 2v_{18} - v_6 - v_{11} - v_{15} - 2v_{17} \\
\frac{d}{dt}[PLN_4^{(p0)}](t) &= v_7 + v_{20} + v_{22} - v_8 - v_{19} - v_{21} \\
\frac{d}{dt}[PLN_4^{(p1)}](t) &= v_9 + v_{24} + v_{26} - v_{10} - v_{23} - v_{25}
\end{aligned}
$$

$$\frac{d}{dt}[PLN_4^{(p2)}](t) = v_{11} + v_{13} + v_{28} + v_{30} - v_{12} - v_{14} - v_{27} - v_{29}$$
$$\frac{d}{dt}[PLN_4^{(p3)}](t) = v_{15} + v_{32} + v_{34} - v_{16} - v_{31} - v_{33}$$
$$\frac{d}{dt}[PLN_4^{(p4)}](t) = v_{17} + v_{36} + v_{38} - v_{18} - v_{35} - v_{37}$$
$$\frac{d}{dt}[PLN_5^{(p0)}](t) = v_{19} + v_{43} + v_{44} - v_{20} - v_{42}$$
$$\frac{d}{dt}[PLN_5^{(p1)}](t) = v_{21} + v_{23} + v_{42} + v_{46} + v_{47} - v_{22} - v_{24} - v_{43} - v_{44} - v_{45}$$
$$\frac{d}{dt}[PLN_5^{(p2)}](t) = v_{25} + v_{27} + v_{45} + v_{49} + v_{50} - v_{26} - v_{28} - v_{46} - v_{47} - v_{48}$$
$$\frac{d}{dt}[PLN_5^{(p3)}](t) = v_{29} + v_{31} + v_{48} + v_{52} + v_{53} - v_{30} - v_{32} - v_{49} - v_{50} - v_{51}$$
$$\frac{d}{dt}[PLN_5^{(p4)}](t) = v_{33} + v_{35} + v_{51} + v_{55} + v_{56} - v_{34} - v_{36} - v_{52} - v_{53} - v_{54}$$
$$\frac{d}{dt}[PLN_5^{(p5)}](t) = v_{37} + v_{54} - v_{38} - v_{55} - v_{56}$$
$$\frac{d}{dt}[Inh\text{-}1^{(p0)}](t) = v_{58} - v_{57}$$
$$\frac{d}{dt}[PP1](t) = v_{60} - v_{59}$$

We do not account for synthesis and degradation of proteins and thus assume the total amount of PLN, inhibitor-1 and PP1 to be conserved. This allows us to determine three species by the following algebraic equations:

$$\begin{aligned}[PLN_1^{(p1)}](t) &= [PLN]_{tot} - [PLN_1^{(p0)}](t) - 2([PLN_2^{(p0)}](t) + [PLN_2^{(p1)}](t) + [PLN_2^{(p2)}](t)) \\ &- 4([PLN_4^{(p0)}](t) + [PLN_4^{(p1)}](t) + [PLN_4^{(p2)}](t) + [PLN_4^{(p3)}](t) + [PLN_4^{(p4)}](t)) \\ &- 5([PLN_5^{(p0)}](t) + [PLN_5^{(p1)}](t) + [PLN_5^{(p2)}](t) + [PLN_5^{(p3)}](t) + [PLN_5^{(p4)}](t) + [PLN_5^{(p5)}](t))\end{aligned}$$

$$[PP1 : Inh\text{-}1^{(p1)}](t) = [PP1]_{tot} - [PP1](t)$$

$$[Inh\text{-}1^{(p1)}](t) = [Inh\text{-}1]_{tot} - [Inh\text{-}1^{(p0)}](t) - [PP1 : Inh\text{-}1^{(p1)}](t),$$

where $[PLN]_{tot}, [PP1]_{tot}$ and $[Inh\text{-}1]_{tot}$ are the conserved total concentrations of PLN, PP1 and inhibitor-1, respectively.

#### 2.3 Reaction rates

**Oligomerization of PLN**

Combinatorial expansion due to phospho-isoforms of oligomeric complexes can lead to thermodynamic inconsistencies if mass action kinetics are applied to the oligomerization reactions as given in a reaction scheme. This can be circumvented by introducing balancing coefficients which account for the effective oligomerization rates regardless of combinatorial effects. We thus modeled oligomerization reactions with balanced mass action kinetics as described in [11].

**Dynamic equilibrium of PLN**

How exactly dynamic equilibrium of PLN works on the molecular level is currently not known, although electrostatic interactions [12, 13] and anionic detergents seem to play an important role. Since electrostatic interactions can increase the association rate constant of an interaction [14], we assumed this to be the case for phosphorylated PLN. We, therefore, introduced a dimensionless control parameter $\omega$, which increases association rate constants of oligomerization reactions in an exponential fashion depending on how many phosphorylated subunits are involved.

**Enzymatic reactions catalysed by PKA, PP1 and PP2A**

Reactions catalysed by PKA, PP2A were modeled using a modified Michaelis-Menten rate law, which accounts for competition between multiple substrates $S_1, ..., S_n$:

$$v_i = \frac{V_{max} S_i}{K_{m_i}(1 + \sum_{j \in J \setminus \{i\}} \frac{S_j}{K_{m_j}}) + S_i},$$

where $J = \{1, ..., n\}$, and $v_i$ describes the rate of consumption of substrate $S_i$ [15]. Since PLN pentamers and monomers are several orders of magnitude more abundant than dimers or tetramers, the possibility of direct dimer or tetramer (de-)phosphorylation can be ignored and was thus not accounted for in this study. To account for the cooperativity observed in PLN pentamer dephosphorylation [16], we assumed that PP1 (since it is the phosphatase responsible for most PLN dephosphorylation [17, 18]) dephosphorylates pentamers by a cooperative mechanism and introduced two dimensionless control parameters $\phi \leq 1$ and $\chi \geq 1$. For pentamers with $n \geq 1$ phosphorylated subunits, we multiplied $k_{cat,PP1:PLN_5}$ with factor $\chi^{n-1}$ and $K_{m,PP1:PLN_5}$ with factor $\chi^{n-1}$. For $\phi < 1$ we thus increase the turnover number for each phosphate group removed from a pentamer, with $k_{cat,PP1:PLN_5}$ being an upper limit. For $\chi > 1$ we thus reduce the Michaelis-constant for each phosphate group removed from a pentamer, with $K_{m,PP1:PLN_5}$ representing highest substrate affinity (lowest $K_m$ value). Such modes of cooperativity can be called *v-type* and *k-type* cooperativity, respectively [19] (Figure S16).

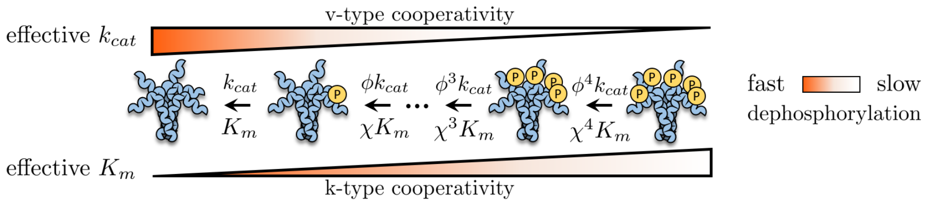

**Figure S16**

Illustration of $PLN_5$ dephosphorylation following the concept of positive $v-/k-type$ cooperativity [19]. For each phosphate group to be removed, the effective $k_{cat}$ or $K_m$ is scaled by factor $\phi < 1$ or $\chi > 1$, respectively.

For simplicity and lack of data suggesting otherwise, we assume $K_{m,PP1:PLN_1} = K_{m,PP1:PLN_5}$ and $k_{cat,PP1:PLN_1} = k_{cat,PP1:PLN_5}$, implying that the kinetic constants of monomer dephosphorylation pose an upper efficiency limit for pentamer dephosphorylation. If one assumes that these pose a lower limit for pentamer dephosphorylation, simulated pentamer phosphorylation occurs slower and at lower steady-state levels than monomers (data not shown), contradicting experimental observations [2]. Potentially, this could be amended by assuming higher $k_{cat}$ and lower $K_m$ values for pentamer phosphorylation by PKA compared to monomers. However, since available kinetic data is not sufficient to distinguish between these possibilities, our current implementation of cooperative pentamer dephosphorylation is one of the simplest and requires the fewest assumptions.

Following the outlined rationale of the chosen rate laws, the complete list of the reaction rates of the model is given by:

**Oligomer association/dissociation rates**

$$v1 = k_1 \cdot [PLN_1^{(p0)}]^2$$
$$v2 = k_2 \cdot [PLN_2^{(p0)}]$$
$$v3 = 2 \cdot k_1 \cdot [PLN_1^{(p0)}] \cdot [PLN_1^{(p1)}]$$
$$v4 = k_2 \cdot [PLN_2^{(p1)}]$$
$$v5 = k_1 \cdot \omega \cdot [PLN_1^{(p1)}]^2$$
$$v6 = k_2 \cdot [PLN_2^{(p2)}]$$
$$v7 = k_3 \cdot [PLN_2^{(p0)}]^2$$
$$v8 = k_4 \cdot [PLN_4^{(p0)}]$$
$$v9 = 2 \cdot k_3 \cdot [PLN_2^{(p0)}] \cdot [PLN_2^{(p1)}]$$
$$v10 = k_4 \cdot [PLN_4^{(p1)}]$$
$$v11 = 2 \cdot k_3 \cdot [PLN_2^{(p0)}] \cdot [PLN_2^{(p2)}]$$
$$v12 = \frac{1}{2} \cdot k_4 \cdot [PLN_4^{(p2)}]$$
$$v13 = k_3 \cdot \omega \cdot [PLN_2^{(p1)}]^2$$
$$v14 = \frac{1}{2} \cdot k_4 \cdot [PLN_4^{(p2)}]$$
$$v15 = 2 \cdot k_3 \cdot \omega^2 \cdot [PLN_2^{(p1)}] \cdot [PLN_2^{(p2)}]$$
$$v16 = k_4 \cdot [PLN_4^{(p3)}]$$
$$v20 = k_6 \cdot [PLN_5^{(p0)}]$$
$$v21 = k_5 \cdot [PLN_1^{(p1)}] \cdot [PLN_4^{(p0)}]$$
$$v22 = \frac{1}{2} \cdot k_6 \cdot [PLN_5^{(p1)}]$$
$$v23 = k_5 \cdot [PLN_1^{(p0)}] \cdot [PLN_4^{(p1)}]$$
$$v24 = \frac{1}{2} \cdot k_6 \cdot [PLN_5^{(p1)}]$$
$$v25 = k_5 \cdot \omega \cdot [PLN_1^{(p1)}] \cdot [PLN_4^{(p1)}]$$
$$v26 = \frac{1}{2} \cdot k_6 \cdot [PLN_5^{(p2)}]$$
$$v27 = k_5 \cdot [PLN_1^{(p0)}] \cdot [PLN_4^{(p2)}]$$
$$v28 = \frac{1}{2} \cdot k_6 \cdot [PLN_5^{(p2)}]$$
$$v29 = k_5 \cdot \omega^2 \cdot [PLN_1^{(p1)}] \cdot [PLN_4^{(p2)}]$$
$$v30 = \frac{1}{2} \cdot k_6 \cdot [PLN_5^{(p3)}]$$
$$v31 = k_5 \cdot [PLN_1^{(p0)}] \cdot [PLN_4^{(p3)}]$$
$$v32 = \frac{1}{2} \cdot k_6 \cdot [PLN_5^{(p3)}]$$
$$v33 = k_5 \cdot \omega^3 \cdot [PLN_1^{(p1)}] \cdot [PLN_4^{(p3)}]$$
$$v34 = \frac{1}{2} \cdot k_6 \cdot [PLN_5^{(p4)}]$$
$$v35 = k_5 \cdot [PLN_1^{(p0)}] \cdot [PLN_4^{(p4)}]$$

$$v17 = k_3 \cdot \omega^3 \cdot [PLN_2^{(p2)}]^2$$
$$v18 = k_4 \cdot [PLN_4^{(p4)}]$$
$$v19 = k_5 \cdot [PLN_1^{(p0)}] \cdot [PLN_4^{(p0)}]$$
$$v36 = \frac{1}{2} \cdot k_6 \cdot [PLN_5^{(p4)}]$$
$$v37 = k_5 \cdot \omega^4 \cdot [PLN_1^{(p1)}] \cdot [PLN_4^{(p4)}]$$
$$v38 = k_6 \cdot [PLN_5^{(p5)}]$$

#### Phosphorylation and dephosphorylation of PLN

$$v39 = \frac{[PKA]k_{cat,PKA:PLN_1}[PLN_1^{(p0)}]}{K_{m,PKA:PLN_1}\big(1 + \sum_{i\in I}(\frac{[PLN_5^{(pi)}]}{K_{m,PKA:PLN_5}}) + \frac{[Inh\text{-}1^{(p0)}]}{K_{m,PKA:Inh\text{-}1}}\big) + [PLN_1^{(p0)}]}$$

$$v40 = \frac{[PP1]k_{cat,PP1:PLN_1}[PLN_1^{(p1)}]}{K_{m,PP1:PLN_1}\big(1 + \sum_{j\in J}\frac{[PLN_5^{(pj)}]}{\chi^{j-1}K_{m,PP1:PLN_5}}\big) + [PLN_1^{(p1)}]}$$

$$v41 = \frac{V_{max,PP2A:PLN_1}[PLN_1^{(p1)}]}{K_{m,PP2A} + [PLN_1^{(p1)}] + \sum_{j\in J}([PLN_5^{(pj)}]) + [Inh\text{-}1^{(p1)}]}$$

$$v42 = \frac{[PKA]k_{cat,PKA:PLN_5}[PLN_5^{(p0)}]}{K_{m,PKA:PLN_5}\big(1 + \frac{[PLN_1^{(p0)}]}{K_{m,PKA:PLN_1}} + \sum_{i\in I\setminus\{0\}}(\frac{[PLN_5^{(pi)}]}{K_{m,PKA:PLN_5}}) + \frac{[Inh\text{-}1^{(p0)}]}{K_{m,PKA:Inh\text{-}1}}\big) + [PLN_5^{(p0)}]}$$

$$v43 = \frac{[PP1]k_{cat,PP1:PLN_5}[PLN_5^{(p1)}]}{K_{m,PP1:PLN_5}\big(1 + \frac{[PLN_1^{(p1)}]}{K_{m,PP1:PLN_1}} + \sum_{j\in J\setminus\{1\}}\frac{[PLN_5^{(pj)}]}{\chi^{j-1}K_{m,PP1:PLN_5}}\big) + [PLN_5^{(p1)}]}$$

$$v44 = \frac{V_{max,PP2A:PLN_5}[PLN_5^{(p1)}]}{K_{m,PP2A} + [PLN_1^{(p1)}] + \sum_{j\in J}([PLN_5^{(pj)}]) + [Inh\text{-}1^{(p1)}]}$$

$$v45 = \frac{[PKA]k_{cat,PKA:PLN_5}[PLN_5^{(p1)}]}{K_{m,PKA:PLN_5}\big(1 + \frac{[PLN_1^{(p0)}]}{K_{m,PKA:PLN_1}} + \sum_{i\in I\setminus\{1\}}(\frac{[PLN_5^{(pi)}]}{K_{m,PKA:PLN_5}}) + \frac{[Inh\text{-}1^{(p0)}]}{K_{m,PKA:Inh\text{-}1}}\big) + [PLN_5^{(p1)}]}$$

$$v46 = \frac{\phi[PP1]k_{cat,PP1:PLN_5}[PLN_5^{(p2)}]}{\chi K_{m,PP1:PLN_5}\big(1 + \frac{[PLN_1^{(p1)}]}{K_{m,PP1:PLN_1}} + \sum_{j\in J\setminus\{2\}}\frac{[PLN_5^{(pj)}]}{\chi^{j-1}K_{m,PP1:PLN_5}}\big) + [PLN_5^{(p2)}]}$$

$$v47 = \frac{V_{max,PP2A:PLN_5}[PLN_5^{(p2)}]}{K_{m,PP2A} + [PLN_1^{(p1)}] + \sum_{j\in J}([PLN_5^{(pj)}]) + [Inh\text{-}1^{(p1)}]}$$

$$v48 = \frac{[PKA]k_{cat,PKA:PLN_5}[PLN_5^{(p2)}]}{K_{m,PKA:PLN_5}\big(1 + \frac{[PLN_1^{(p0)}]}{K_{m,PKA:PLN_1}} + \sum_{i\in I\setminus\{2\}}(\frac{[PLN_5^{(pi)}]}{K_{m,PKA:PLN_5}}) + \frac{[Inh\text{-}1^{(p0)}]}{K_{m,PKA:Inh\text{-}1}}\big) + [PLN_5^{(p2)}]}$$

$$v49 = \frac{\phi^2[PP1]k_{cat,PP1:PLN_5}[PLN_5^{(p3)}]}{\chi^2 K_{m,PP1:PLN_5}\big(1 + \frac{[PLN_1^{(p1)}]}{K_{m,PP1:PLN_1}} + \sum_{j\in J\setminus\{3\}}\frac{[PLN_5^{(pj)}]}{\chi^{j-1}K_{m,PP1:PLN_5}}\big) + [PLN_5^{(p3)}]}$$

$$v50 = \frac{V_{max,PP2A:PLN_5}[PLN_5^{(p3)}]}{K_{m,PP2A} + [PLN_1^{(p1)}] + \sum_{j\in J}([PLN_5^{(pj)}]) + [Inh\text{-}1^{(p1)}]}$$

$$v51 = \frac{[PKA]k_{cat,PKA:PLN_5}[PLN_5^{(p3)}]}{K_{m,PKA:PLN_5}\big(1 + \frac{[PLN_1^{(p0)}]}{K_{m,PKA:PLN_1}} + \sum_{i\in I\setminus\{3\}}(\frac{[PLN_5^{(pi)}]}{K_{m,PKA:PLN_5}}) + \frac{[Inh\text{-}1^{(p0)}]}{K_{m,PKA:Inh\text{-}1}}\big) + [PLN_5^{(p3)}]}$$

$$v52 = \frac{\phi^3[PP1]k_{cat,PP1:PLN_5}[PLN_5^{(p4)}]}{\chi^3 K_{m,PP1:PLN_5}\big(1 + \frac{[PLN_1^{(p1)}]}{K_{m,PP1:PLN_1}} + \sum_{j\in J\setminus\{4\}}\frac{[PLN_5^{(pj)}]}{\chi^{j-1}K_{m,PP1:PLN_5}}\big) + [PLN_5^{(p4)}]}$$

$$v53 = \frac{V_{max,PP2A:PLN_5}[PLN_5^{(p4)}]}{K_{m,PP2A} + [PLN_1^{(p1)}] + \sum_{j\in J}([PLN_5^{(pj)}]) + [Inh\text{-}1^{(p1)}]}$$

$$v54 = \frac{[PKA]k_{cat,PKA:PLN_5}[PLN_5^{(p4)}]}{K_{m,PKA:PLN_5}\big(1 + \frac{[PLN_1^{(p0)}]}{K_{m,PKA:PLN_1}} + \sum_{i\in I\setminus\{4\}}(\frac{[PLN_5^{(pi)}]}{K_{m,PKA:PLN_5}}) + \frac{[Inh\text{-}1^{(p0)}]}{K_{m,PKA:Inh\text{-}1}}\big) + [PLN_5^{(p4)}]}$$

$$v55 = \frac{\phi^4[PP1]k_{cat,PP1:PLN_5}[PLN_5^{(p5)}]}{\chi^4 K_{m,PP1:PLN_5}\big(1 + \frac{[PLN_1^{(p1)}]}{K_{m,PP1:PLN_1}} + \sum_{j\in J\setminus\{5\}}\frac{[PLN_5^{(pj)}]}{\chi^{j-1}K_{m,PP1:PLN_5}}\big) + [PLN_5^{(p5)}]}$$

$$v56 = \frac{V_{max,PP2A:PLN_5}[PLN_5^{(p5)}]}{K_{m,PP2A} + [PLN_1^{(p1)}] + \sum_{j\in J}([PLN_5^{(pj)}]) + [Inh\text{-}1^{(p1)}]}$$

**Reactions involving inhibitor-1**

$$v57 = \frac{[PKA]k_{cat,PKA:Inh\text{-}1}[Inh\text{-}1^{(p0)}]}{K_{m,PKA:Inh\text{-}1}\big(1 + \sum_{i\in I}(\frac{[PLN_5^{(pi)}]}{K_{m,PKA:PLN_5}}) + \frac{[PLN_1^{(p0)}]}{K_{m,PKA:PLN_1}}\big) + [Inh\text{-}1^{(p0)}]}$$

$$v58 = \frac{V_{max,PP2A:Inh\text{-}1}[Inh\text{-}1^{(p1)}]}{K_{m,PP2A} + [PLN_1^{(p1)}] + \sum_{j\in J}([PLN_5^{(pj)}]) + [Inh\text{-}1^{(p1)}]}$$

$$v59 = k_7[Inh\text{-}1^{(p1)}][PP1]$$

$$v60 = k_8[PP1 : Inh\text{-}1^{(p1)}]$$

#### 2.4 Model parameters and initial conditions

A mathematical model needs parameters and initial conditions to make useful predictions. In order to obtain realistic parameters values and protein concentrations we searched the literature and previously published models related to calcium handling and $\beta$-adrenergic signaling in cardiomyocytes, as well as phospholamban, PKA, PP1 and inhibitor-1.

**Table S1**
Review initial conditions / protein concentrations

| Protein concentration | Value | Source | Comments |
|---|---|---|---|
| $[PLN]_{tot}$ | ≈ 250 $\mu$M | this study | cf. Figure S14 |
| | 106 $\mu$M | [20] | |
| | >50 $\mu$M | [21, 22] | |
| | 38 $\mu$M | [23] | |
| $[PKA]$ | 0.59 $\mu$M | [20] | |
| | 0.5176 $\mu$M | [24] | |
| | 0.48 $\mu$M | [25] | |
| $[PP1]_{tot}$ | 0.89 $\mu$M | [20, 25] | |
| | 0.2 $\mu$M | [24] | |
| | 0.5 $\mu$M* | [26] | (skeletal muscle) |
| $[Inh\text{-}1]_{tot}$ | 0.3 $\mu$M | [20] | |
| | 0.08543 $\mu$M | [24] | |

* Calculated by the number of protein molecules per cell given in the reference and the rule of thumb that in an "average" eukaryotic cell, 1000 molecules of a protein roughly correspond to a cellular concentration of 1 nM ([27]; BioNumbers BNID 104519).

**Table S2**
Review kinetic and equilibrium constants

| Parameter | Value | Source | Comments |
|---|---|---|---|
| $k_{cat,PKA:PLN}$ | 54 $s^{-1}$ | [20] | ($k_{cat}$ for PLN phosphorylation by PKA) |
| | 21 $s^{-1}$ | [22] | |
| | 23.4 $s^{-1}$ | [28] | $PLN_{1-20}$, no lipid environment |
| | 22.3 – 25 $s^{-1}$ | [29] | $PLN_{1-19}$, different environments |
| | 13 – 21.9 $s^{-1}$ | [29] | AFA-PLN, different environments |
| $K_{m,PKA:PLN}$ | 21 $\mu$M | [20] | ($K_m$ for PLN phosphorylation by PKA) |
| | 12.5 $\mu$M | [22] | |
| | 93.3 $\mu$M | [28] | $PLN_{1-20}$, no lipid environment |
| | 36.4 – 90.1 $\mu$M | [29] | $PLN_{1-19}$, different environments |
| | 47.6 – 238.1*$\mu$M | [29] | AFA-PLN, different environments |

* Not included in calculation of median value for default parameter set.

Review kinetic and equilibrium constants (continued)

| Parameter | Value | Source | Comments |
|---|---|---|---|
| $k_{cat,PP1:PLN}$ | 8.5 $s^{-1}$ | [20] | ($k_{cat}$ for PLN phosphorylation by PKA) |
| $K_{m,PP1:PLN}$ | 7 $\mu$M | [20] | ($K_m$ for PLN dephosphorylation by PP1) |
| $V_{max,PP2A:PLN}$ | 0.708 $\mu$M $s^{-1}$ | [17] | guesstimate$^{\#}$, ($V_{max}$ for PLN dephosphorylation by PP2A) |
| $V_{max,PP2A:Inh\text{-}1}$ | 14 $\mu$M $s^{-1}$ | [20] | ($V_{max}$ for inhibitor-1 dephosphorylation by PP2A) |
| $k_{cat,PKA:Inh\text{-}1}$ | 60 $s^{-1}$ | [20] | ($k_{cat}$ for inhibitor-1 phosphorylation by PKA) |
| $K_{m,PKA:Inh\text{-}1}$ | 1 $\mu$M | [20] | ($K_m$ for inhibitor-1 phosphorylation by PKA) |
| $K_{m,PP2A}$ | 1 $\mu$M | [20] | ($K_m$ for inhibitor-1 dephosphorylation by PP2A) |
| $K_{d,PP1:Inh\text{-}1^{(p1)}}$ | 1 nM | [20] | ($K_d$ for complex of PP1 and phosphorylated inhibitor-1) |

### Macdougall *et al.* reported that PP1 accounts for 70-90% and PP2A for the remaining dephosphorylation activity towards phospholamban (and to a small extend PP2C, too) [17]. We thus assumed that PP2A dephosphorylation accounts for approximately 20% of the total dephosphorylation activity towards phospholamban observed by Macdougall *et al.*. We furthermore assumed the same $K_m$ value as for inhibitor-1 dephosphorylation.

**Parameter $\omega$ (dynamic equilibrium of PLN)**
Since individual oligomerization steps and the influence of phosphorylation on their rate constants have not been studied before, we used our own data and those from Hou *et al.* [13] to obtain a first estimate for $\omega$ as described in subsubsection 3.1.2.

| Parameter | Value | Source | Comments |
|---|---|---|---|
| $\omega$ | $\approx 1.066$ | this study | based on data from Figure S19C |
| | $\approx 1.044$ | [13] | see subsubsection 3.1.2 for calculation |

It is important to note, however, that the extend to which this effect can be observed varies considerably. Wittmann *et al.*, for instance, observed no significant increase in pentamerization upon S16 phosphorylation in forskolin stimulated HEK293 cells [2]. Cornea *et al.*, in contrast, found a near complete pentamerization of Ser16-phosphorylated, recombinant PLN in DOPC lipid bilayers using electron paramagnetic resonance spectroscopy [12]. The effect observed by Hou *et al.* [13], could be an underestimation given that phospho-mimetic mutations are not always faithful experimental models for real phosphorylation [30]. Moreover, the lipid environment, too, plays an important role for both pentamerization [31] and the dynamic equilibrium effect (this study). Taken together, these considerations make it difficult to get a precise estimate from the data so far. We thus decided to opt for a 'guesstimate' of $\omega = 1.25$, which is higher than the estimates based on our or Hou *et al.*'s data, but still lower than what one would expect for the near complete pentamerization observed by Cornea *et al.*

**Cooperative dephosphorylation of pentamers**
To the best of our knowledge, no kinetic parameters for the (de-)phosphorylation of pentameric phospholamban have been determined so far. In the absence of better evidence we mostly assumed the parameters to be the same as for monomeric phospholamban. An exception to this is the dephosphorylation of pentameric PLN by PP1. In order to implement the positive cooperativity of dephosphorylation reported by Li *et al.* [16], we assumed a pronounced v-type cooperativity of $\phi = 0.2$ and the absence of k-type cooperativity, i.e. $\chi = 1$.

**Set of default parameters and initial conditions**
Based on these reviews and considerations we composed a default parameter set, which was used for all simulations and analyses unless specified otherwise in the simulation protocols (section 4). Oligomerization parameter values are from the best fit parameter set of mass action model 2 calibrated with our experimental data (see subsection 2.1).

**Table S3**
Default parameter values and initial conditions

| Parameter/IC | Value | Comments |
|---|---|---|
| $[PLN]_{tot}$ | 250 $\mu$M | |
| $[PKA]$ | 0.59 $\mu$M | |
| $[PP1]_{tot}$ | 0.89 $\mu$M | free $[PP1](t=0) = [PP1]_{tot}$ unless stated otherwise |
| $[Inh\text{-}1]_{tot}$ | 0.3 $\mu$M | $[Inh\text{-}1^{(p0)}](t=0) = [Inh\text{-}1]_{tot}$ unless stated otherwise |
| $k_1$ | 144724 mol s$^{-1}$ | (rate constant for dimer formation; best fit parameter set) |
| $k_2$ | 5518.45 s$^{-1}$ | (rate constant for dimer dissociation; best fit parameter set) |
| $k_3$ | 72672.8 mol s$^{-1}$ | (rate constant for tetramer formation; best fit parameter set) |
| $k_4$ | 1.20089 s$^{-1}$ | (rate constant for tetramer dissociation; best fit parameter set) |
| $k_5$ | 1.83 $\times 10^8$ mol s$^{-1}$ | (rate constant for pentamer formation; best fit parameter set) |
| $k_6$ | 0.13275 s$^{-1}$ | (rate constant for pentamer dissociation; best fit parameter set) |
| $k_7$ | $5\times10^4$ mol s$^{-1}$ | (association rate constant for binding of phosphorylated inhibitor-1 to PP1; matched with $k_8$ to a $K_d$ of 1 nM) |
| $k_8$ | $5\times10^{-5}$ s$^{-1}$ | (dissociation rate constant of complex between phosphorylated inhibitor-1 and PP1; matched with $k_7$ to a $K_d$ of 1 nM) |
| $\omega$ | 1.25 | (increased oligomerization upon PLN phosphorylation) |
| $k_{cat,PKA:PLN_1}$ | 23 s$^{-1}$ | ($k_{cat}$ for PLN$_1$ phosphorylation by PKA; median of reviewed values) |
| $K_{m,PKA:PLN_1}$ | 42 $\mu$M | ($K_m$ for PLN$_1$ phosphorylation by PKA; median of reviewed values) |
| $k_{cat,PKA:PLN_5}$ | 23 s$^{-1}$ | ($k_{cat}$ for PLN$_5$ phosphorylation by PKA; assumed equal to monomers) |
| $K_{m,PKA:PLN_5}$ | 42 $\mu$M | ($K_m$ for PLN$_5$ phosphorylation by PKA; assumed equal to monomers) |
| $k_{cat,PP1:PLN_1}$ | 8.5 s$^{-1}$ | ($k_{cat}$ for PLN$_1$ dephosphorylation by PP1) |
| $K_{m,PP1:PLN_1}$ | 7 $\mu$M | ($K_m$ for PLN$_1$ dephosphorylation by PP1) |
| $k_{cat,PP1:PLN_5}$ | 8.5 s$^{-1}$ | (baseline $k_{cat}$ for PLN$_5$ dephosphorylation by PP1) |
| $K_{m,PP1:PLN_5}$ | 7 $\mu$M | (baseline $K_m$ for PLN$_5$ dephosphorylation by PP1) |
| $V_{max,PP2A:PLN}$ | 0.708 $\mu$M s$^{-1}$ | ($V_{max}$ for PLN dephosphorylation by PP2A) |
| $V_{max,PP2A:Inh\text{-}1}$ | 14 $\mu$M s$^{-1}$ | ($V_{max}$ for inhibitor-1 dephosphorylation by PP2A) |
| $k_{cat,PKA:Inh\text{-}1}$ | 60 s$^{-1}$ | ($k_{cat}$ for inhibitor-1 phosphorylation by PKA) |
| $K_{m,PKA:Inh\text{-}1}$ | 1 $\mu$M | ($K_m$ for inhibitor-1 phosphorylation by PKA) |
| $K_{m,PP2A}$ | 1 $\mu$M | ($K_m$ for inhibitor-1 and PLN dephosphorylation by PP2A) |
| $K_{d,PP1:Inh\text{-}1^{(p1)}}$ | 1 nM | ($K_d$ for complex of PP1 and phosphorylated inhibitor-1) |
| $\phi$ | 0.2 | (v-type cooperativity for PLN$_5$ dephosphorylation by PP1) |
| $\chi$ | 1 | (k-type cooperativity for PLN$_5$ dephosphorylation by PP1) |

### 3 Methods

#### 3.1 Computational procedures

##### 3.1.1 Model implementation and software

All models were implemented as MATLAB® (v2019b) scripts for numerical simulation and analysis. Simulations were performed with the ode23s integrator on an Asus® laptop PC with Intel® Core™i7-7500U CPU @ 2.70GHz, 2904 Mhz, 2 Core(s) processor, 8GB RAM running under Microsoft® Windows 10 OS. Simulation protocols describing the used parameter values and initial conditions for each simulation shown in the figures of the main text (where deviating from the default parameter set) can be found in section 4. The model code is accessible in the BioModels database under: https://www.ebi.ac.uk/biomodels/MODEL2011110001.

##### 3.1.2 Parameter estimation and model selection

Parameter estimation for association and dissociation rate constants of the oligomerization reactions in our mass-action kinetics models was performed in COPASI (v4.26) using the genetic algorithm with 250 generations and a population size of 40. Parameters were constrained by the detailed balance relationships described subsection 2.1. Furthermore, association rate constants were constrained to lie between $10^3 \text{mol}\cdot\text{s}^{-1}$ and $10^9 \text{mol}\cdot\text{s}^{-1}$, whereas dissociation constants were constrained to lie between $10^{-5}\text{s}^{-1}$ and $10^6\text{s}^{-1}$. For each model variant, 30 independent parameter estimation runs with randomized initial values parameters were performed. For model selection, the AIC was calculated as described in [32] by $AIC = 2 \cdot k + n \cdot (ln(\frac{SSE}{n}) + 1)$, where $k$ is the number of model parameters, $n$ is the number of experimental observations and $SSE$ is the sum of weighted squared errors for the best fit parameter set of a model. The AIC score balances how well a model fits the data versus model complexity in number of parameter values and penalizes deviations from experimental data and high number of model parameters. The lower the AIC score, the better. If two models fit the data equally well, the model with fewer parameters is rewarded a lower AIC score because it has less unnecessary complexity.

Estimation of parameter $\omega$ to our own data was performed by assuming that the increased pentamerization shown in figure Figure S19 reflects the steady state situation and by using the MATLAB® function `nlinfit` to fit the parameter to this data.

Calculation of parameter $\omega$ based on the data from Hou *et al.* [13] was done as follows. The apparent monomer-pentamer equilibrium in a monomer-dimer-tetramer-pentamer model such as ours can be characterized by the apparent dissociation constant:

$$K_{d\ 1,5}^{app} = \frac{[PLN_1]^5}{[PLN_5]}.$$

Since we have

$$\begin{aligned} [PLN_5] &= K_{4,5}[PLN_4][PLN_1] \\ &= K_{2,4}K_{4,5}[PLN_2]^2[PLN_1] \\ &= K_{1,2}^2 K_{2,4}K_{4,5}[PLN_1]^5, \end{aligned}$$

we can write

$$K_{d\ 1,5}^{app} = \frac{[PLN_1]^5}{[PLN_5]} = \frac{1}{K_{1,2}^2 K_{2,4} K_{4,5}} = \frac{k_2^2 k_4 k_6}{k_1^2 k_3 k_5},$$

where in the last step the equilibrium constants were substituted with their constitute rate constants. For completely phosphorylated PLN, the derivation of the apparent dissociation constant is identical except for factoring in $\omega$ into the individual equilibria:

$$K_{d\ 1,5}^{app,pPLN} = \frac{[PLN_1^{(p1)}]^5}{[PLN_5^{(p5)}]} = \left(\frac{k_2}{\omega k_1}\right)^2 \left(\frac{k_4}{\omega^3 k_3}\right)\left(\frac{k_6}{\omega^4 k_5}\right) = \frac{1}{\omega^9}\frac{k_2^2 k_4 k_6}{k_1^2 k_3 k_5}.$$

Using the apparent $K_d$ values from Hou *et al.* (given in arbitrary units) for S16A PLN (unphosphorylated) and S16E PLN (mimicking phosphorylated PLN) we arrive at:

$$\frac{K_{d\ 1,5}^{app}}{K_{d\ 1,5}^{app,pPLN}} = \frac{2.2}{1.5} = \frac{\frac{k_2^2 k_4 k_6}{k_1^2 k_3 k_5}}{\frac{1}{\omega^9}\frac{k_2^2 k_4 k_6}{k_1^2 k_3 k_5}} = \omega^9 \Leftrightarrow \omega = \sqrt[9]{\frac{2.2}{1.5}} \approx 1.044$$

##### 3.1.3 Bifurcation diagrams

Bifurcation diagrams were generated using a custom algorithm described previously [11] to make the implementation of bifurcation analysis into the simulation/analysis pipeline easier (e.g. for sensitivity analysis of the bistable range). The algorithm iteratively identifies/approximates the unstable steady states of bistable systems numerically, but does not detect other types of bifurcations (e.g. Hopf-bifurcations). For more details, please refer to: https://www.ebi.ac.uk/biomodels/MODEL1910220002.

##### 3.1.4 Local sensitivity analysis

Local sensitivity analyses was performed as described in [33]. We analysed two steady states which were chosen to be sufficiently remote from the saddle node bifurcations SN1 and SN2 of the model: a low relative phosphorylation level ($[PKA] = 0.13\mu M$), and a high relative phosphorylation level ($[PKA] = 0.25\mu M$). Relative sensitivities of (relative) PLN monomer phosphorylation at steady state upon perturbing the nominal value of each parameter $p$ by $dp = 1\%$ at a time were calculated as $s_r(p) = \frac{drPLN_1^{(p1)}/rPLN_1^{(p1)}}{dp/p}$, where $rPLN_1^{(p1)} = \frac{[PLN_1^{(p1)}]}{[PLN_1^{(p0)}]+[PLN_1^{(p1)}]}$. For bifurcation sensitivity analysis, relative sensitivities were determined in a similar fashion, but instead of focussing on the steady state concentration of relative PLN monomer phosphorylation, relative sensitivities of the bistable range (|SN1 - SN2|) were calculated. In order to obtain well detectable changes in relative sensitivities with a limited granularity of the bifurcation algorithm, a perturbation of $dp = 10\%$ was chosen.

##### 3.1.5 Probing the multi-dimensional parameterspace for its influence on bistability

In order to probe how bistability depends on the multi-dimensional parameter space of our model, we decided to perform an analysis developed by [34]. First, we used the review of parameter values and protein concentrations to guide the construction of ranges of physiologically plausible values in which the true values are likely to lie. As we could not get our model to be executed by the python implementation provided in [34], we decided to implement the analysis in the MATLAB® code of our model. However, as our model equations are quite stiff for some combinations of parameters, we first performed a model reduction and replaced one ODE for an algebraic expression in order to speed up the numerical solution of the model. Due to ease of implementation, a quasi-steady-state (QSS) approximation of $PP1$ is particularly suited to replace an ODE with an algebraic expression. Although this can affect the model dynamics (and thereby eliminate e.g. the response delay of the inhibitor-1 FFL), we used the reduced model only for steady state analyses for which the QSS assumption is (trivially) fulfilled [33]. At steady state we find that:

$$\frac{d}{dt}[PP1](t) = k_8[Inh\text{-}1^{(p1)} : PP1](t) - k_7[Inh\text{-}1^{(p1)}](t)[PP1](t) = 0$$

Assuming that $PP1$ is quasi at steady state compared to the other processes in the reaction network, we denote its concentration in the reduced model by $[\tilde{PP}1]$. From subsection 2.2 we can identify the following conservation laws:

$$[PP1]_{tot} = [PP1](t) + [PP1 : Inh\text{-}1^{(p1)}](t)$$
$$[Inh\text{-}1]_{tot} = [Inh\text{-}1^{(p0)}](t) + [Inh\text{-}1^{(p1)}](t) + [PP1 : Inh\text{-}1^{(p1)}](t)$$

Applying these to the first equation and solving the resulting quadratic equation for $[\tilde{PP}1]$ using the pq-formula gives (for physical reasons only the positive solution is given):

$$[\tilde{PP}1](t) = -\left(\frac{k_8 + k_7([Inh\text{-}1]_{tot} - [Inh\text{-}1^{(p0)}](t) - [PP1]_{tot})}{2k_7}\right) + \sqrt{\frac{(k_8 + k_7([Inh\text{-}1]_{tot} - [Inh\text{-}1^{(p0)}](t) - [PP1]_{tot})^2)}{4k_7^2} - \frac{k_8}{k_7}[PP1]_{tot}}$$

Having eliminated the ODE for PP1 with this algebraic expression, we observed no deviations in the steady state behavior but a marked performance gain.

For the intended analysis, parameters and initial conditions were sampled randomly from a uniform distribution determined by a range of physiologically plausible values. Using the reduced model, two simulations with either completely unphosphorylated or completely phosphorylated PLN (both monomeric and pentameric) as initial conditions were performed with simulation endpoint $t_{end} = 5 \times 10^5 s$. To test whether the model reached steady state, the algorithm checked whether monomer concentration was stable for at least three time points before reaching the end of the simulation. A parameter set was considered bistable if there was a relative difference in phosphorylation levels of >10%. Parameter sets without a difference in phosphorylation levels or with a difference of <10% were considered monostable since we reasoned that marginal differences in steady state phosphorylation might not be of physiological relevance (although technically still bistable). Parameter sets were normalized to the maximum value of the respective parameter range and visualized using parallel coordinate plots.

##### 3.1.6 Frequency response analysis

Frequency response analysis was performed as described in [33, 35]. First, the system's jacobian matrix

$$\mathbf{A} = \frac{\partial \mathbf{f}}{\partial \mathbf{x}} = \begin{pmatrix} \frac{\partial f_1}{\partial [PLN_1^{(p0)}]} & \frac{\partial f_1}{\partial [PLN_2^{(p0)}]} & \cdots & \frac{\partial f_1}{\partial [PP1]} \\ \frac{\partial f_2}{\partial [PLN_1^{(p0)}]} & \frac{\partial f_2}{\partial [PLN_2^{(p0)}]} & \cdots & \frac{\partial f_2}{\partial [PP1]} \\ \vdots & \vdots & \ddots & \vdots \\ \frac{\partial f_{17}}{\partial [PLN_1^{(p0)}]} & \frac{\partial f_{17}}{\partial [PLN_2^{(p0)}]} & \cdots & \frac{\partial f_{17}}{\partial [PP1]} \end{pmatrix},$$

where $f_1, ..., f_{17}$ ($\mathbf{f}$ in vector notation) denotes the right hand sides of the ODEs in subsection 2.2 as a function of the model variables $[PLN_1^{(p0)}], ..., [PP1]$ ($\mathbf{x}$ in vector notation) in the specified order, was calculated symbolically using MATLAB®. For the purpose of the frequency response analysis, the model input $u$ was defined as $[PKA]$ and the output $h$ as relative monomer phosphorylation $\frac{[PLN_1^{(p1)}]}{[PLN_1^{(p0)}]+[PLN_1^{(p1)}]}$. Linearized input and output maps $\mathbf{B}$ and $\mathbf{C}$ were calculated as $\mathbf{B} = \frac{\partial \mathbf{f}}{\partial u}$ and $\mathbf{C} = \frac{\partial \mathbf{h}}{\partial \mathbf{x}}$, respectively. The feed-through term $D$ as $\frac{\partial \mathbf{h}}{\partial u}$. After evaluating $\mathbf{A}$,$\mathbf{B}$,$\mathbf{C}$ and $D$ at a nominal operating point, a linearized input-output system was created using the MATLAB® function `ss2tf`. The linearized input-output system was subsequently used for the evaluation of the system bandwidth and for creating Bode-diagrams using the MATLAB® functions `bandwidth` and `bode`, respectively.

#### 3.2 Experimental procedures

##### Reconstitution of PLN in detergent micelles

PLN (human wild-type and monomeric 'AFA'-mutant C36A, C41F, C46A ) was purchased as a synthetic peptide from Pepscan (Lelystad, Netherlands). Successful synthesis of the peptide, as indicated by the correct molecular weight, was confirmed by Pepscan via mass-spectrometry (Figure S17). For further analysis, 1 mg of lyophilized peptide was resolubilized in 1 ml of buffer for 2 hours at room temperature under gentle overhead agitation (10 rpm) and subsequently centrifuged in a bench-top centrifuge for 10 min at >10000 ×g, 4°C to get rid of insoluble residual material. For experiments involving PLN phosphorylation, we tested various detergents in a buffer with physiological ionic strength and pH and found that Triton X-100, which is a milder detergent than the harsh ionic SDS, is excellent at solubilizing PLN at rather low concentrations and allows for effective pentamerization as well as rapid phosphorylation by PKA. We thus used the following Triton X-100 based buffer (TBB) for all experiments involving PLN phosphorylation: 50 mM TRIS-HCl pH 7.5, 100 mM NaCl, 10% (v/v) glycerol, 0.5% (v/v) Triton X-100, 2 mM DTT.

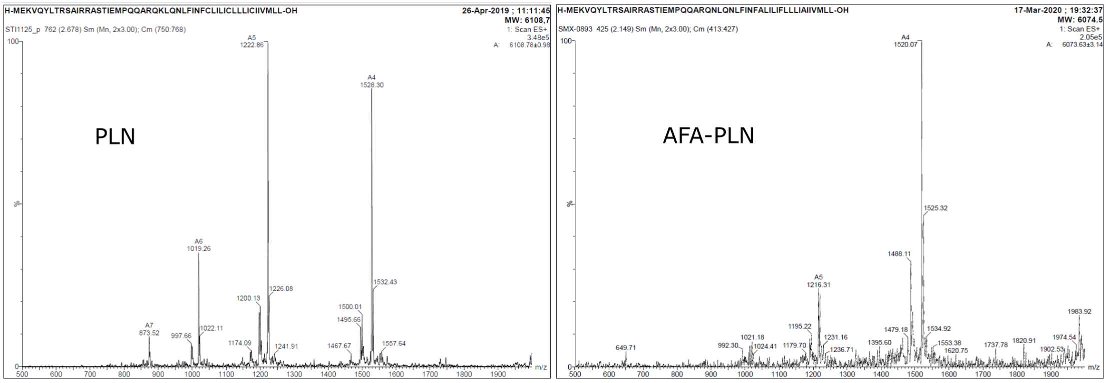

**Figure S17**
ES-MS spectra of human wild-type PLN (left) and AFA-PLN (right).

Interestingly, although showing less pentamerization than wild-type PLN, AFA-PLN was not entirely monomeric when kept on ice or even at room temperature. However, short heating for 15 min at 50°C was sufficient to dissociate all pentamers into monomers (Figure S18). Although some smearing of band was visible after SDS-PAGE, no precipitation in solution was visible even after 45 min at 50°C and AFA-PLN still appears to be an excellent substrate for PKA after heating, which indicates that heating PLN for 15 min at 50°C does not seriously denature the protein. Once dissociated, no significant re-oligomerization was observed after incubation at 25°C for 2 hours (the time frame for our phosphorylation experiments) or after snap-freezing and quick thawing.

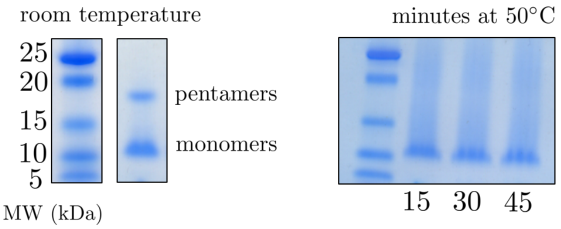

**Figure S18**
SDS-PAGE analysis of the oligomeric status of AFA-PLN at different temperatures and incubation times. 0.33 mg/ml protein concentration, loaded 2 µl per lane.

#### Semi-native SDS-PAGE

PLN forms stable pentamers even in harsh SDS sample buffer which only dissociate upon sample boiling. As SDS sample buffer is likely to interfere with less stable proteins (e.g. enzymes), all phosphorylation reactions were performed in TBB and analysed using semi-native SDS-PAGE by directly applying the native samples (i.e. phosphorylated or unphosphorylated PLN in TBB) into the wells of the gels (4–20% Mini-PROTEAN® TGX™ Precast Protein Gels, Bio-Rad, cat.no. 4561096) without addition of other buffers or boiling. All samples contained 10% glycerol to allow samples to settle into the wells and to avoid mixing with running buffer.

#### Oriole staining and quantification of absolute concentrations

Since conventional Coomassie staining of protein gels only has limited sensitivity and more sensitive silver-staining protocols are typically not suitable for quantitative purposes, we used Oriole™ Fluorescent Gel Stain for detection and quantification of PLN (Bio-Rad, cat. no. 161-0496). Staining was performed for 90 min according to the manufacturer's instructions. Gels were imaged on a ChemiDocTM XRS+ imaging system (Bio-Rad) and data were quantified using the ImageLab v6.0 software (Bio-Rad). The method shows a wide linear range for PLN and is neither affected by oligomerization or phosphorylation status of PLN (Figure S19). Since the Oriole™ signal is not affected by oligomeric state, the signal for an oligomeric species is directly proportional to the number protomers in a complex. Knowing the total PLN concentration of a sample thus allows to calculate the absolute concentrations of monomers and pentamers by $[\mathrm{PLN}_x] = \frac{F^{Oriole}_{PLN_x} \cdot [\mathrm{PLN}_{tot}]}{x}$, where $F^{Oriole}_{PLN_x}$ is the fraction which $\mathrm{PLN}_x$ contributes to the total Oriole™ signal. In either SDS-sample buffer or TBB we observed no other PLN forms than monomers and pentamers and occasionally a weak band for dimers at higher concentrations. Since the dimer band was not well demarked and negligible compared to the monomer and pentamer bands we did not quantify dimers.

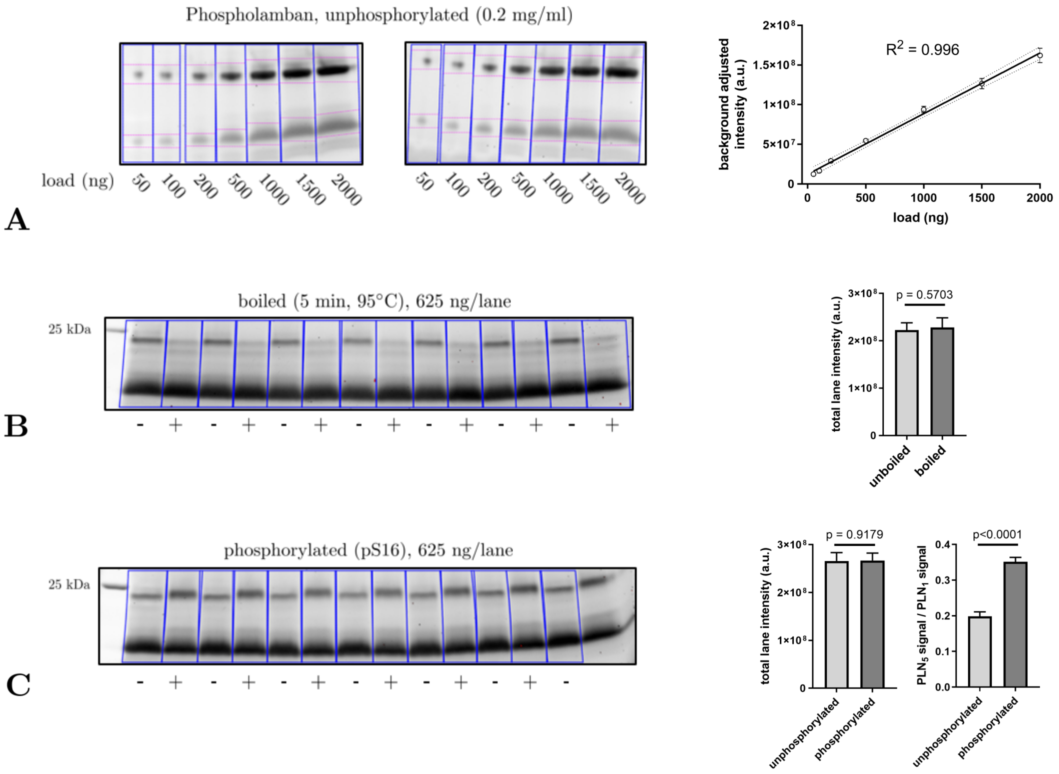

**Figure S19**

Evaluation of Oriole™ Fluorescent Gel Stain for quantification of PLN. (A) the signal of total PLN (monomers + pentamers) in TBB shows excellent linearity for the probed range (50-2000 ng). (B) 1 mg/ml of boiled/unboiled PLN in TBB was diluted to 0.05 mg/ml in 1x Laemmli buffer and loaded to the gel. Quantification of total lane intensities suggests that the oligomeric state of PLN does not influence the signal intensity. (C) 1 mg/ml of phosphorylated/unphosphorylated PLN in TBB was diluted to 0.05 mg/ml in 1x SDS-sample buffer and loaded to the gel. Quantification of total lane intensities suggests that phosphorylation state of PLN does not influence the signal intensity. The increased pentamer/monomer signal ratio suggests that S16 phosphorylation leads to increased pentamerization in mixed SDS/Triton X-100 micelles.

#### Dilution experiments

Samples were diluted in TBB or SDS with end concentrations ranging from ≈ 0.96 mg/ml (undiluted) to 5 μg/ml in a final volume of 40µl. After dilution, samples were left to pre-equilibrate for 45 min at room temperature before being incubated in a PCR machine for 75 min at 37°C with heated lid at 50°C to avoid evaporation. To avoid disturbing the pentamerization equilibrium by sample cooling, semi-native SDS-PAGE was performed using SDS-running buffer pre-warmed to 37°C. Total protein amount for each well was within the determined linear range.

For production of phosphorylated PLN, TBB samples were supplemented with 250 µM ATP, 5 mM $MgCl_2$ and 3 U/µl PKA (Sigma-Aldrich, cat. no. 539576; a kind gift of Dr. Thomas Kampourakis). Phosphorylation was allowed to proceed for 16 hours at 4°C and given another hour at room temperature to approach completion. To avoid differences due to ionic strength, precipitation or evaporation, samples for unphosphorylated PLN in TBB were treated accordingly, but without addition of PKA.

##### Dissociation time-course experiments

To determine pentamer dissociation dynamics, 40µl of 0.96 mg/ml PLN in TBB supplemented with 250 µM ATP and 5 mM $MgCl_2$ were pre-equilibrated for 30 min at RT followed by 30 min at 37°C. Samples were diluted 20-fold with 37°C pre-warmed TBB and incubated for up to 15 min at 37°C. To simultaneously determine the oligomeric status at different time points, dilutions were started in replicates at 1.5, 7.5 and 15 min before semi-native SDS-page (performed with 37°C pre-warmed running buffer). Samples were processed and loaded to the gel using a multichannel pipette to minimize sample processing time.

##### *In vitro* phosphorylation time-courses of PLN by PKA

To determine the competitive effect of pentamers on monomer phosphorylation, we compared the phosphorylation dynamics of wild-type PLN and AFA-PLN. To ensure complete dissociation into monomers, AFA-PLN was heated to 50 °C for 15 min followed by 10 min incubation at 25 °C immediately before the experiment. Phosphorylation reactions were performed at 25°C in a volume of 150 µl TBB supplemented with 250 µM ATP, 5 mM $MgCl_2$ and 6.25 U/µl (≈ 7.7 nM) PKA (Sigma-Aldrich, cat. no. 539576; a kind gift of Dr. Thomas Kampourakis) with an end concentration of 0.962 mg/ml ($\approx 157 \mu M$) wild-type PLN or 0.32 mg/ml AFA-PLN (at which there is an equimolar monomer concentration of ≈ 52 µM between wild-type PLN and AFA-PLN). Samples (8 µl) were taken at 0, 5, 15, 30, 60, 90 and 120 min, snap-frozen and stored in liquid nitrogen until separation by semi-native SDS-PAGE (1.5 µl/well for wild-type PLN, 2 µl/well for AFA-PLN). Phosphorylation was detected using Pro-Q$^{TM}$ Diamond staining (Thermo Fisher, cat. no. P33300) according to the manufacturer's instructions (1 h staining step), followed by Quick Coomassie Staining (Generon, cat. no. NB-45-00078-1L) to visualize total protein amount. Gels were imaged on a ChemiDocTM XRS+ imaging system (Bio-Rad) and data were quantified using the ImageLab v6.0 software (Bio-Rad). The phosphorylation signal of each experiment was corrected by total protein amount as given by the Coomassie signal and by subtracting the background signal at 0 min. Relative phosphorylation levels were calculated by fitting progress curves of wild-type pentamers and AFA-$PLN_1$ to a hyperbola ($Y(t) = S_{max} * t/(K + t)$ and dividing the corrected phosphorylation signal by $S_{max}$ (the signal expected for complete phosphorylation). Since phosphorylation of wild-type monomers was still linear during the probed reaction period (preventing reliable fitting to a hyperbola), relative phosphorylation was calculated by using $S_{max}$ from wild-type pentamers scaled by $r = 52\mu M/(157 - 52)\mu M$, the amount of phosphorylation sites in monomers relative to the amount of phosphorylation sites in pentamers at the given total concentration of PLN.

##### PKA-dependent phosphorylation of PLN in transfected HEK293 cells

cDNA of PLN was cloned into pcDNA3 expression vector and expressed in HEK293 cells as described previously [2]. PLN-expressing cells were then treated for 40 min with forskolin (Santa Cruz) in Dulbecco's Modified Eagle's Medium (DMEM) at 37°C at different concentrations to induce PKA-dependent phosphorylation of PLN. Cells were washed with PBS before mechanical lysis in PBS containing protease inhibitors and phosphatase inhibitors. Lysates were centrifuged for 20 min at >10,000 g at 4°C and supernatants were used for western blot analysis. Equal amounts of protein (Pierce® BCA Assay Kit, ThermoScientific) were separated on 15% polyacrylamide gels and transferred to PVDF membranes (Immobilon®-P, Millipore) before overnight incubation with primary antibodies in TBS-T / 5% milk (10 mM Tris, 150 mM NaCl, 0.1 % Tween 20, pH 7.6, 5% milk) at 4°C. Antigen detection was performed by chemiluminescence using secondary antibodies coupled to horseradish peroxidase (abcam) and Luminata Forte Western HRP substrate (Millipore). The following antibodies were used for detection of proteins: anti-PLN (A1, Badrilla, 1:5,000 dilution), anti-phospho-PLN (S16, Badrilla, 1:5,000 dilution). Data were quantified using the ImageLab v6.0 software (Bio-Rad).

#### 3.3 Statistical analysis

Statistical comparison between two experimental groups was performed in GraphPad Prism (v8.2) using the two-tailed unpaired T-test, corrected for multiple comparisons where appropriate. Sample sizes are given in the respective figures. A p-value $< 0.05$ was considered to be statistically significant. Significance levels were encoded as follows: ns = not significant, * $p < 0.05$, ** $p < 0.01$, *** $p < 0.001$, **** $p < 0.0001$.

### 4 Appendix: simulation protocols

To enable reproducibility of our simulations in other software environments, this appendix elaborates the simulation protocols, initial conditions and parameter values underlying the presented simulations in the main text. Figure numbers refer to the main text. For specifications of the simulations from the supplementary material, please refer to the published code.

- **Figure 3A, left**: First, 250$\mu$M $[PLN]_{tot}$ (unphosphorylated) were simulated until oligomerization reached equilibrium in the absence of any other species (no inhibitor-1, no phosphatases, no PKA). Subsequently, [PKA] = 0.1$\mu$M and a 300 s phosphorylation time-course of PLN species was simulated in the absence of phosphatases and inhibitor-1. Other initial conditions and parameters as listed in Table S3.
- **Figure 3A, right**: PKA was set to 0 $\mu$M and final PLN species concentrations from Figure 3A (left) were used as initial conditions for PLN. Free PP1 was set 0.1 $\mu$M and a 250 s dephosphorylation time-course was simulated. Inhibitor-1 was absent. PP2A and other initial conditions and parameters as listed in Table S3.
- **Figure 3B**: red and blue lines are the same data as in figure 3A replotted as relative monomer and pentamer phosphorylation, respectively. For the simulation in the absence of pentamers (dotted purple line), the simulations described for Figure 3A were repeated with rate constants $k_1 = k_2 = k_3 = k_4 = k_5 = k_6 = 0$ (to prevent oligomerization) and initial PLN concentration equimolar to the monomer concentration of $[PLN]_{tot} = 250$ $\mu$M at oligomerization equilibrium ($\approx 58.2\mu$M).
- **Figure 3E**: First, 250$\mu$M $[PLN]_{tot}$ (unphosphorylated) were simulated until oligomerization reached equilibrium in the absence of any other species (no inhibitor-1, no phosphatases, no PKA). Subsequently, PKA was set to 0.35 $\mu$M and a 350s phosphorylation time-course of PLN species was simulated the presence of phosphatases and inhibitor-1 (concentrations as listed in Table S3). Subsequently, PKA was set to 0 $\mu$M and dephosphorylation was simulated for 250 s. The whole simulation was performed three times with different rate constants for binding of inhibitor-1 to PP1: $k_7 = 1/2/4 \times 10^4$ mol s$^{-1}$ and $k_8 = 1/2/4 \times 10^{-5}$ s$^{-1}$, respectively. Finally, the simulation was repeated in the absence of inhibitor-1 but with PP1 reduced by the fraction of PP1 bound to inhibitor-1 at steady state from previous simulations (to ensure the same steady state is approached). Other initial conditions and parameters as listed in Table S3.
- **Figure 3F**: (e1-e3) $[PLN_1](t{=}0) = [PLN]_{tot} = 55$ $\mu$M, $k_1 = k_2 = k_3 = k_4 = k_5 = k_6 = 0$, $V_{max,PP2A:PLN} = V_{max,PP2A:Inh-1} = 0$, $k_{cat,PKA:Inh-1} = 0.25$ s$^{-1}$, $K_{m,PKA:Inh-1}$= 14$\mu$M, $t_{end} = 2000s$. (e1) [PKA] = 10 nM, [PP1](t=0) = $[PP1]_{tot}$ = 1$\mu$M, [Inh-1$^{(p0)}$](t=0) = $[Inh\text{-}1]_{tot}$ = 10$\mu$M. (e2) [PKA] = 10 nM, [PP1](t=0) = $[PP1]_{tot}$ = 0$\mu$M, [Inh-1$^{(p0)}$](t=0) = $[Inh\text{-}1]_{tot}$ = 0$\mu$M. (e3) [PKA] = 10 nM, [PP1](t=0) = $[PP1]_{tot}$ = 1$\mu$M, [Inh-1$^{(p0)}$](t=0) = $[Inh\text{-}1]_{tot}$ = 0$\mu$M.
- **Figure 4A**: First, 250$\mu$M $[PLN]_{tot}$ (unphosphorylated) were simulated until oligomerization reached equilibrium in the absence of any other species (no inhibitor-1, no phosphatases, no PKA). Next, ten simulations with 0-100% (in steps of 10%) of monomeric and pentameric PLN from the first step being phosphorylated at t=0 were run for 2000s at [PKA] = 0.21$\mu$M and other initial conditions and parameters as listed in Table S3.
- **Figure 4B**: Bifurcation plots were calculated as described in subsubsection 3.1.3. PKA was chosen as bifurcation parameter in the range of 0.1 to 0.6 $\mu$M, $t_{end} = 500000$ s and 30 iterations were allowed for direct identification of unstable steady states before the algorithm switches to approximative identification. When the unstable steady state was approximated, the first 30 integration steps were ignored in order to prevent a bias resulting from smaller integration steps at the beginning of the simulations.
- **Figure 4D**: As described in subsubsection 3.1.4.
- **Figure 4E**: As described in subsubsection 3.1.5.
- **Figure 5A**: First, 250$\mu$M $[PLN]_{tot}$ (unphosphorylated) were simulated until oligomerization reached equilibrium in the absence of any other species (no inhibitor-1, no phosphatases, no PKA). Second, except for [PKA], all species and parameters were set to their default value as listed in Table S3. Next, a series of short bursts of [PKA] = 0.59$\mu$M, each 45 s apart, was simulated and burst duration was increased for each simulation (1/3.3/10s).
- **Figure 5B**: As in Figure 5A but in absence of inhibitor-1 and with PP1 reduced by the fraction of PP1 bound to inhibitor-1 at steady state from previous simulations (to ensure the same steady state is approached).

- **Figure 5C**: As in Figure 5A but with rate constants $k_1 = k_2 = k_3 = k_4 = k_5 = k_6 = 0$ (to prevent oligomerization) and an initial PLN concentration equimolar to the monomer concentration of $[\text{PLN}]_{tot} = 250\ \mu$M at oligomerization equilibrium ($\approx 58.2\mu$M).
- **Figure 5D**: combined settings of Figure 5B and Figure 5C.
- **Figure 5E**: First, the model was simulated to steady state with [PKA] = $0.2\mu$M and other initial conditions and parameters as listed in Table S3. From the steady state, we simulated the model for 512 short time intervals of 0.1 s length each, where for the i-th time interval, we set $[\text{PKA}](\text{i}) = 0.2\mu M + 0.05 \cdot 0.2\mu M \cdot sin(\frac{(i-1)}{20}) \pm rnd() \cdot 0.075 \cdot 0.2\mu M$, where the last summand represents a fast random noise component.
- **Figure 5F**: First, the model was simulated to steady state with [PKA] = $0.2\mu$M and other initial conditions and parameters as listed in Table S3. Frequency response analysis for the reached steady state was carried out as described in subsubsection 3.1.6. For frequency response analysis in the absence of pentamers or inhibitor-1, simulation until steady state was modified as described for Figure 5B and 5C.
- **Figure 5G**: The parameters of the parameter set yielding ultrasensitivity without bistability which deviate from Table S3 are:

| **Parameter/IC** | **Value** | **Comments** |
|---|---|---|
| $[PP1]_{tot}$ | 0.67 $\mu$M | free $[PP1](t=0) = [PP1]_{tot}$ unless stated otherwise |
| $K_{m,PKA:PLN_1}$ | 55 $\mu$M | ($K_m$ for $\text{PLN}_1$ phosphorylation by PKA) |
| $K_{m,PKA:PLN_5}$ | 42.5 $\mu$M | ($K_m$ for $\text{PLN}_5$ phosphorylation by PKA) |
| $k_{cat,PP1:PLN_5}$ | 9 $s^{-1}$ | (baseline $k_{cat}$ for $\text{PLN}_5$ dephosphorylation by PP1) |
| $K_{m,PP1:PLN_5}$ | 4 $\mu$M | (baseline $K_m$ for $\text{PLN}_5$ dephosphorylation by PP1) |
| $\phi$ | 0.79 | (v-type cooperativity for $\text{PLN}_5$ dephosphorylation by PP1) |
| $\chi$ | 1.03 | (k-type cooperativity for $\text{PLN}_5$ dephosphorylation by PP1) |

  Bifurcation diagrams were calculated as described for Figure 4B. The response to fluctuations in PKA concentration was determined as follows: First, the model was simulated to steady state with [PKA] = $0.226\mu$M and other initial conditions and parameters as listed in Table S3 or as given in the table above for the parameter set leading to ultrasensitivity without bistability. From the steady state, we simulated the model for 100 time intervals of 60 s length each, where for the i-th time interval, we set $[\text{PKA}](\text{i}) = 0.226\mu M \pm rnd() \cdot 0.25 \cdot 0.226\mu M$.
- **Figure 5H,I**: Noise-landscapes were calculated as follows. First, the model was simulated to steady state at a given baseline PKA concentration $[\text{PKA}]_{BL}$ and other initial conditions and parameters as listed in Table S3 or as given in the table above for the parameter set leading to ultrasensitivity without bistability. From the steady state, we simulated the model for 150 time intervals of 60 s length each, where for the i-th time interval, we set $[\text{PKA}](\text{i}) = [PKA_{bl}] \pm rnd() \cdot n_f \cdot [PKA_{bl}]$, where $n_f$ is the input noise level. From these simulations, the coefficient of variation ($\text{CV} = \frac{\sigma}{\mu}$) of relative PLN monomer phosphorylation was determined, where $\sigma$ and $\mu$ denote the standard deviation and mean of the data, respectively. This process was repeated iteratively and the resulting CVs were visualized on a surface plot as function of the indicated ranges of $[\text{PKA}]_{BL}$ and input noise levels. For the relative noise landscape shown in Figure 5I, CVs resulting from the parameter set for ultrasensitive without bistability were divided by the CVs resulting from the parameter set for bistability.
- **Figure 5J**: The model was simulated to steady state at a given baseline PKA concentration $[\text{PKA}]_{BL}$, where $[\text{PKA}]_{BL} = 0.208\ \mu$M for the parameter set exhibiting bistability or $[\text{PKA}]_{BL} = 0.225\ \mu$M for the parameter set resulting in ultrasensitivity without bistability. Other initial conditions and parameters as listed in Table S3 or as given in the table above for the parameter set leading to ultrasensitivity without bistability. From the steady state, we simulated the model for 1000 time intervals of 60 s length each, where for the i-th time interval, we set $[\text{PKA}](\text{i}) = [PKA_{bl}] \pm rnd() \cdot 0.0625\mu M$. A switching event between low and high phosphorylation states was counted when relative monomer phosphorylation changed from $\leq 0.25$ to $\geq 0.4$ or vice versa.
